# Effect of Age and Sex on BCAA Metabolism in Mice

**DOI:** 10.1101/2025.10.30.685560

**Authors:** Gagandeep Mann, Stephen Mora, Olasunkanmi A J Adegoke

## Abstract

Increased plasma levels of branched-chain amino acids (BCAA) have been implicated in insulin resistance. This condition worsens with age, but plasma BCAA levels are downregulated in old men. The effect of age and/ sex on BCAA metabolism has been rarely studied. Thus, the objective of this study was to analyze how age and sex affect BCAA levels and their metabolism in mouse tissues.

Male and female young (4-month) and old (18-months) CD2F1 mice were used. BCAA levels and relevant BCAA metabolic enzymes abundance/activity in plasma, muscle, liver, adipose tissue, and heart were analyzed.

Old males exhibited greater plasma BCAA concentrations compared to young males and old females, but plasma branched-chain ketoacid (BCKA) levels were lower in old mice in both sexes. Intracellular BCAA levels were lower in skeletal muscle and heart from old mice independent of sex. There was an age-sex interaction in the lever in that reduced total BCAA was seen only in old male animals. In adipose tissue, total BCKA levels were higher in old male animals. Skeletal muscle abundance of the BCAA transporter LAT1 was reduced in old females compared to young females. In the liver, female mice had higher levels of LAT1 than male mice independent of age, while total BCKD was reduced in old females compared to old males. Pp2Cm levels were reduced in old animals independent of sex.

In conclusion, while there were some changes in plasma and tissue BCAA/BCKA levels in response to age/sex, such changes were largely not consistent with changes in tissue BCAA catabolic enzyme abundance/activity. This suggests that protein levels of BCAA catabolic enzymes are preserved in aging in healthy animals and likely only become dysregulated in disease states.

## Introduction

Branched-chain amino acids (BCAA; leucine, valine, isoleucine) are essential amino acids required in a human diet. BCAA stimulate muscle protein synthesis and regulate body weight and glucose homeostasis ^(1)^. Their catabolism begins with transamination by branched-chain aminotransferase 2 (BCAT2), producing branched-chain α-ketoacids (BCKA; alpha-ketoisocaproic acid (KIC), alpha-keto-beta methylvaleric acid (KMV), and alpha-ketoisovaleric acid (KIV), respectively from leucine, isoleucine and valine). Skeletal muscles are the main site of this step. The BCKA are then predominantly catabolized in the liver as the rate limiting enzyme responsible for BCKA catabolism, branched-chain ketoacid dehydrogenase (BCKD), is mostly abundant and active in the liver ^(2)^.

Circulating levels of BCAA are altered in various diseases. In chronic renal failure ^(3,4)^ and liver cirrhosis ^(5,6)^, there is a reduction in plasma BCAA levels, while in maple syrup urine disease ^(7,8)^, and insulin resistant states like obesity and type 2 diabetes mellitus (T2DM), plasma levels are elevated ^(9–11)^. In patients with chronic renal failure, there is an increase in BCKD complex abundance ^(12)^ and BCAA oxidation ^(13)^ in skeletal muscle, which could contribute to reduced muscle and plasma BCAA levels. In liver cirrhosis, decreased BCAA plasma levels may be a product of increased BCKD complex activity ^(14)^. In insulin resistant states, the abundance and activity of BCAA catabolic enzymes are altered. BCAT2 and BCKDHβ mRNA levels are decreased in diabetic patients, which could explain the increase in BCAA levels ^(15)^. Also, BCKD activity is reduced in mouse models of insulin-resistant states like T2DM and obesity ^(16)^. Additionally, BCAA and their metabolites appear causative in inducing insulin resistance ^(17–21)^. Consistent with this, others have examined how targeting BCAA catabolism can potentially improve insulin sensitivity ^(15,16,22,23)^, emphasizing the link between the two pathways.

There is some evidence to link aging with alterations to BCAA metabolism. Old men exhibit lower circulating BCAA levels ^(24,25)^. This is interesting, as insulin resistance is more prevalent in older individuals ^(26)^, and BCAA levels are higher in insulin resistance ^(10,11)^. BCAA concentrations ^(24,25)^ and protein intake ^(27)^ are reduced with aging, which could play a factor, but the activity and the abundance of enzymes involved in BCAA metabolism could also play a role in regulating BCAA levels and insulin sensitivity during aging. To our knowledge, only one study has looked at the effect of age on BCAA metabolic enzymes, and they saw that BCAT2 and BCKDβ protein levels were reduced in adipose tissue of old mice (18-month) compared to young mice (3-month). These old mice also exhibited greater plasma BCAA ^(28)^.

The sex hormone estrogen inhibits the BCKD complex by activating BCKD kinase (BDK), a negative regulator of BCKD, thus decreasing BCAA catabolism ^(29,30)^. Also, plasma BCAA concentrations are higher in males compared to females ^(31)^, and males have greater leucine oxidation compared to women following endurance exercise ^(32)^. However, not much is known about the effect of sex on BCAA levels and BCAA catabolic enzyme activity and abundance in relevant tissues. Thus, since BCAA are implicated in many diseases, and age is a risk factor of many of those diseases, we studied how age affects BCAA levels and enzymes related to BCAA metabolism in various tissues in male and female mice.

## Materials and Methods

### Ethics Statement

All animal experiments were approved by the York University Animal Care Committee and were conducted in line with the guidelines of the Canadian Council on Animal Care (Protocol #2020-09).

### Animals

Fifteen male and 15 female 8-week-old CD2F1 were purchased from Charles River (Senneville, Quebec, Canada) or Envigo Laboratories (Haslett, Michigan, United States). This study was done in parallel with other studies in which we are examining the effects of cancer and chemotherapy on BCAA metabolism. We are using the CD2F1 strain in those studies because these mice are not as refractory to tumors compared to the C57 strains ^(33–37)^. Since BCAA metabolism has not been studied in CD2F1 and we intend to relate data from this study to those from our cancer/chemotherapy studies, we chose to use the CD2F1 strain. Mice were acclimatized and housed in the vivarium with free access to food and water. Animals were handled 2–3 times per week to reduce stress of handling them on the day of the experiment. Young (4-month) female and male mice (n=7 for each) and old (18-month) female and male mice (n=8 for each) were used. We considered 18-month mice as “old”, consistent with other aging studies looking at changes in plasma BCAA ^(28,38,39)^. For females, estrous tracking was done to ensure that female mice were in the same stage of their estrous cycle. Animal handling and tissue harvesting were conducted in a careful manner to minimize stress and pain as described ^(40)^.

### Reagents

Anti gamma tubulin antibody (#T6557), amino acid standard (#AAS-18), O-phthalaldehyde (OPA, #P1378), 1,2-diamino-4,5-methylenedioxybenzene (DMB, #66807), valine (#V0513), leupeptin (#L2884), coenzyme A (CoA, #4282), NAD+ (#N0632), thiamine (#T1270), magnesium chloride (#7786-30-3), and protease (#P8430) and phosphatase #P5726) inhibitor cocktails were purchased from Sigma Aldrich (Oakville, Ontario, Canada). Insulin (Humulin R, DIN#00586714) was purchased from Eli Lilly Canada Inc (Toronto, Ontario, Canada). Glucometer (#71675) and glucose strips (#71681) were purchased from AlphaTrak, (Cleveland, Ohio, United States). Antibody against BDK (#PA5-31455), Pierce bicinchoninic acid (BCA) protein assay kit (#23225) and fetal bovine serum (FBS, #12483-020) were purchased from Thermo Fisher Canada (Burlington, Ontario Canada). Antibodies against phosphorylated (ph) BCKDH E1α (Ser293, #40368), BCKD-E1α (#90198), SDHA (#11998), as well as horseradish peroxidase (HRP) conjugated anti rabbit (#7074) and anti mouse (#7076) secondary antibodies were purchased from Cell Signaling Technology (Danvers, Massachusetts, United States). Antibody against BCAT2 (#16417-1-AP) and pp2Cm (14573-1-AP) were purchased from ProteinTech (Rosemont, Illinois, United). U-^14^C labelled valine (#ARC0678) was purchased from American Radiolabeled Chemicals (St. Louis, Missouri, United States of America). Chemiluminescence substrate (#WBKLS0500) was purchased from Millipore (Etobicoke, Ontario, Canada). Potassium phosphate monobasic (#PPM302.1), and potassium phosphate dibasic (#PPD555.1) were bought from BioShop (Burlington, ON, Canada).

### Insulin Tolerance Test

Following a 6-h food deprivation, blood was collected through the saphenous vein and used to determine basal glucose levels with the use of AlphaTrak glucose strips and glucometer. Then an intraperitoneal insulin (Humulin R) injection was administered (0.75 U/kg body weight). As done for basal glucose readings, blood samples were collected post insulin (5-120 min) injection and used to determine blood glucose levels as done previously ^(40)^. Animals were euthanized via cervical dislocation. Following sacrifice, plasma, skeletal muscle, adipose tissue, liver, and heart were collected.

### BCKD Activity Assay

Protocol was adapted from White et al ^(23)^. About 30 mg of frozen tibialis anterior, adipose tissue, heart, or liver was crushed in liquid nitrogen and homogenized (Bio-Gen PRO200 Homogenizer, Connecticut, United States) in 250 µL of ice-cold buffer 1 (30 mM KPI, 3 mM EDTA, 5 mM DTT, 1 mM valine, 3% FBS, 5% Triton X-100, 1 µM leupeptin). Homogenates were centrifuged at 10,000 g for 10 minutes. Then, 50 μL of the resultant supernatant was added to an Eppendorf tube with 300 μL of buffer 2 (50 mM HEPES, 30 mM KPI, 0.4 mM CoA, 3 mM NAD+, 5% FBS, 2 mM thiamine, 2 mM magnesium chloride and 7.8 µM ^14^C-labelled valine). This Eppendorf tube now with the homogenate in buffer 1 mixed with buffer 2 had a wick taped to the inside surface of the lid. This wick was prior impregnated with 60 μL of 1 M NaOH. Each Eppendorf tube was capped and tape-sealed tight, before being placed in an incubator at 37°C for 30 min. The radiolabeled ^14^CO_2_ released from the BCKD reaction was captured by the NaOH on the wick, forming NaH^14^CO3 on the wick. The wick was counted in a liquid scintillation counter to measure BCKD activity (amount of CO^2^ released from the BCKD reaction). BCKD activity was calculated by dividing the radioactivity counts by the specific activity of reaction mix (7.8 µM ^14^C-labelled valine in buffer 2) to obtain pmol of CO_2_ released from the oxidative decarboxylation reaction. This value was then divided by the protein concentration in each well to obtain pmol of CO_2_ released from the oxidative decarboxylation reaction per μg protein. Protein content was determined by the Pierce BCA Protein Assay Kit. Absorbance readings at 550nm were obtained using KC4 plate reader software (Bio-Tek Instruments Inc., Winooski, Vermont, United States). Protein concentrations were derived from a standard curve using BSA as the standard.

### Western Blotting

Sections of the tibialis anterior, adipose tissue, liver and heart, were weighed and homogenized in 7 volumes of homogenization buffer (in mM: 20 mM HEPES pH 7.4, 2 mM EGTA, 50 mM NaF, 100 mM KCl, 0.2 mM EDTA, 50 mM glycerol phosphate) supplemented with 1 mM DTT, 1 mM benzamidine, 0.5 mM sodium vanadate, and 10 μl/ml of each of protease and phosphatase inhibitor cocktails. This homogenate was then centrifuged for 3 min at 1000 *g* and 4°C. The supernatant was collected and centrifuged 30 min at 10000 g and 4°C. The Pierce BCA Protein Assay Kit was used to determine protein concentration as mentioned above under BCKD assay. A standard curve was then used to estimate the volume needed to load 40 μg of protein into one well of a polyacrylamide gel. Proteins in homogenate were separated on 10% SDS polyacrylamide gel electrophoresis (SDS PAGE) followed by transfer onto polyvinylidene difluoride (PVDF) membranes (0.2 μm pore size). Membranes were incubated with a ponceau S dye to confirm proteins were transferred. Membranes with ponceau S dye were imaged with BioRAD ChemiDoc XRS+ and quantified with Image Lab software as a loading control. The ponceau S dye was then washed off 3 X 5 minutes washes in Tris Buffered Saline with Tween (TBST). Membranes were then incubated for 1 h in 5% non-fat milk in TBST at room temperature to block non-specific antigen binding. After, they were rinsed 3 X 5 minutes each with TBST at room temperature and then incubated with the primary antibody of interest overnight at 4°C. Following the overnight incubation in primary antibody, membranes were rinsed 3 X 5 minutes each with TBST and then incubated in a secondary antibody (diluted in 5% milk with TBST) for 3 h at room temperature. Secondary antibody (HRP-conjugated anti-rabbit IgG) was diluted in 1:2000-10000 depending on the primary antibody. After incubation with the secondary antibody, membranes were rinsed 3 X 5 minutes each with TBST before HRP chemical luminescent substrate was applied to them. BioRAD ChemiDoc XRS+ was used for signal visualization and the images were quantified with Image Lab software (version 8).

### Amino Acid Concentrations

Plasma samples were centrifuged at 15 min at 3000 g and 4°C. Plasma supernatants and supernatants from tibialis anterior, liver, and heart samples that were homogenized and centrifuged as explained above were diluted in a 1:2:1:8 ratio (sample: potassium phosphate buffer: 0.1 N hydrochloric acid: HPLC grade water, respectively). Diluted samples werepre column derivatized with a 1:1 ratio of sample to OPA (29.28 mM). They were injected into a YMC Triart C18 column (C18, 1.9 μm, 75 × 3.0 mm; YMC America, Allentown, PA, USA) fitted onto an ultra high pressure liquid chromatography system (Nexera X2, Shimadzu, Kyoto, Japan) that was connected to a fluorescence detector (Shimadzu, Kyoto, Japan; excitation: 340 nm; emission: 455 nm). Amino acids were eluted with a gradient solution derived from mobile phase A (20 mM potassium phosphate buffer (6.5 pH)) and mobile phase B (45% HPLC-grade acetonitrile, 40% HPLC-grade methanol and 15% HPLC-grade water) at a flow rate of 0.8 ml/min. We used a gradient of 5%–100% of mobile phase B over 21 min. Amino acid concentrations were calculated using amino acid standard curves. To validate the protocol, each of the BCAA was run through the column to establish elution time for each amino acid. Once this was established, the area of the peak was taken for each point on the standard curve. The amino acid standard curve was used to convert peak areas to amino acid concentrations. For muscle, liver, and heart, data were normalized to total protein concentration of the respective samples using the Pierce BCA protein assay kit as described above.

### BCKA Concentrations

Protocol was adapted from Fujiwara et al ^(41)^. Plasma supernatants and supernatants from homogenized muscle, liver, and heart samples were diluted in a 1:2:1:8 ratio (sample: potassium phosphate buffer: HPLC-grade water/homogenization buffer: HPLC-grade water, respectively). Diluted samples were treated with a 1:1 ratio of a DMB solution (DMB, 13.32 mM), sodium sulfite (38.88 mM), 2-mercaptethanol (1 M), HCl (0.696 M) in ddH_2_O). Once samples were treated with DMB, this solution was heated at 85°C for 45 minutes. After, they were cooled on ice for at least 5 min. They were then injected into an Inertsil ODS-4 column (2 μM, 100 × 2.1 mm; GL Sciences, Torrance, CA, USA) fitted onto an ultra high pressure liquid chromatography system (Nexera X2, Shimadzu, Kyoto, Japan) that was connected to a fluorescence detector (Shimadzu, Kyoto, Japan; excitation: 367 nm; emission: 446 nm). Mobile phases were (A) HPLC-grade methanol/HPLC-grade H_2_O (30/70, v/v) and (B) HPLC-grade methanol. Gradient elution was performed as follows: 0 min 0% B, 3.33 min 0%B, 5 min 50%B, 17.34 min 50%B. The flow rate was 0.2 mL/min, and the column temperature was maintained at 40°C. To validate the protocol, each of the BCKA was run through the column to establish elution times for each BCKA. Once this was done, peak area was taken for each point on the standard curve. The BCKA standard curve was used to convert peak areas in samples to BCKA concentrations. For muscle, liver, and heart, data were normalized to total protein concentration of the respective samples using the Pierce BCA protein assay kit as described above.

### Data Presentation and Statistical Analysis

All immunoblot analyses were quantified, and Ponceau S stains were used for total protein normalization. Phosphorylated BCKD was expressed relative to total BCKD. BCKD activity data is presented as pmol/μg of protein. Data are presented as mean ± SEM. Two-way analysis of variance (ANOVA) was used and Tukey’s post hoc tests were done to measure statistically significant differences among means. Significance was determined as *p* < 0.05. Statistical analyses were performed using GraphPad Prism 10 software (GraphPad, Massachusetts, United States).

## Results

### Old males have increased plasma BCAA levels, while plasma BCKA are reduced in old age

There was an interaction of age and sex on BCAA in that old male mice had greater total plasma BCAA compared to young male mice. The values in old male were also higher than in old female mice (Fig 1A, p<0.05). Plasma leucine, isoleucine, and valine levels were higher in old male mice compared to old female mice (Fig 1A, p<0.05). Old male mice had greater plasma isoleucine (Fig 1A, p<0.05). In female, there was no statistically significant effect of age on these measures.

**Fig 1:**
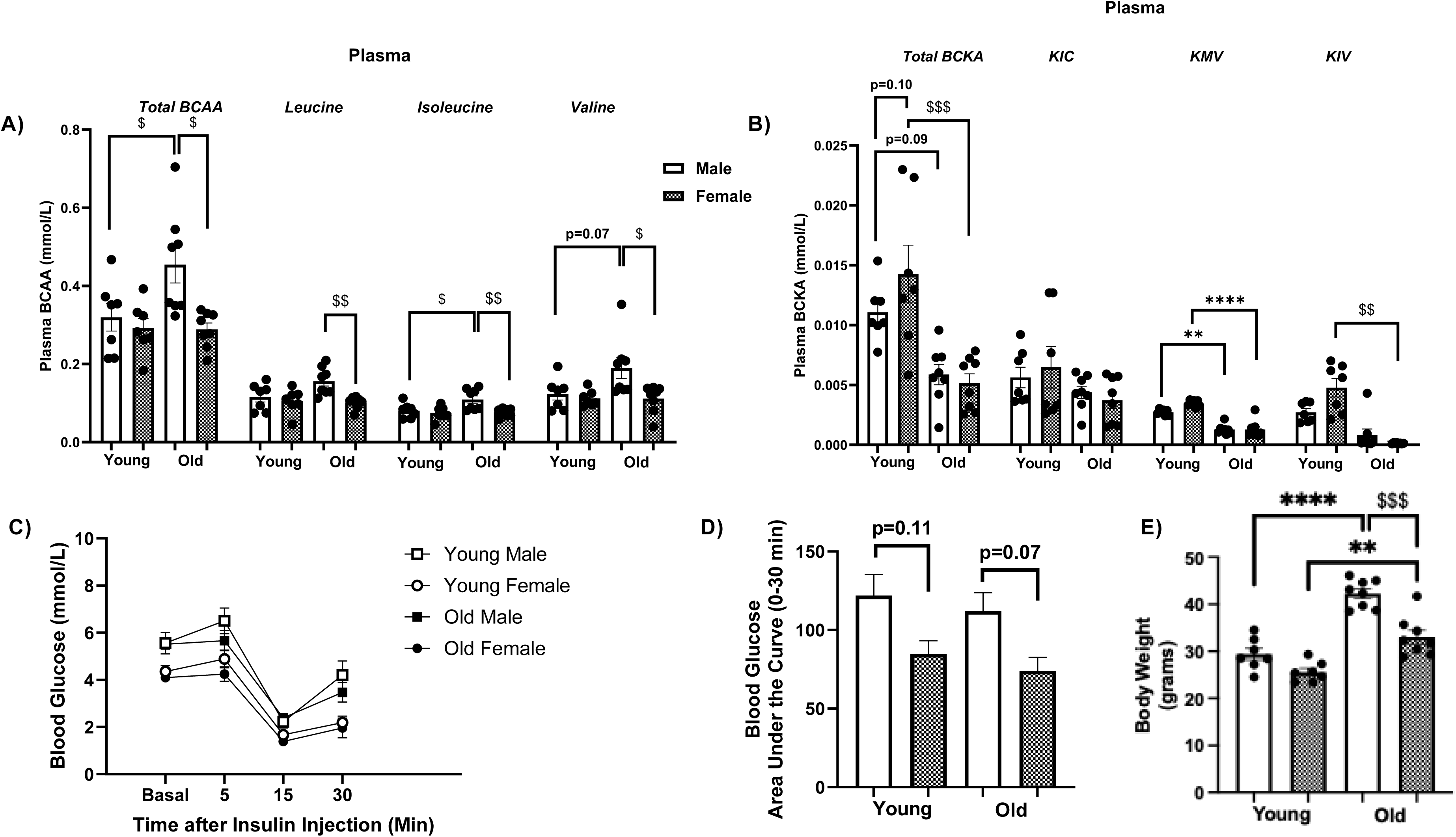
Plasma total BCKA levels reduced in old age, but old male mice exhibit higher plasma BCAA levels. Muscle samples from young (n=7) and old (n=8) mice were analyzed by HPLC to measure plasma total BCAA, leucine, isoleucine, valine (A), and total BCKA, KIV, KIC and KMV (B) concentrations. Blood glucose (basal) was taken right before insulin injection. Insulin was injected and then blood glucose was measured at different timepoints (C). Area under the curve of the blood glucose curve was analyzed (D). The final body weights of the mice are shown (E). Data are means ± SEM. Effect of age: ** p<0.01, **** p<0.0001. Effect of age-sex interaction: $ p<0.05, $$ p<0.01, $$$ p<0.001. # p<0.05, young female vs old female. ^ p<0.05, young male vs old male. Comparisons between groups are shown by lines. For example, in Fig 1A, total plasma BCAA levels in old male are compared to values in young male. Similarly, values in old male are compared to values in old female, and so on.

There was an interaction of age and sex on total plasma BCKA, as old female mice had lower total plasma BCKA than young female mice (Fig 1B, p<0.001). Plasma KIV levels were reduced in old female mice compared to young female mice (Fig 1B, p<0.01). There was an age effect on plasma KMV levels, being lower in old animals (Fig 1B, p<0.01). There was no sex effect on plasma BCKA. In general, the changes in circulating total/individual BCKA in old male are the reciprocal of changes in total/individual BCAA.

Glutamate is produced in the first step of BCAA oxidation ^(42)^. There was no age or sex effect on plasma glutamate levels (Supplementary Fig S1A). There was an age effect on plasma serine and alanine levels, with the older animals having higher levels (Fig S1A). There was an age-sex interaction effect on plasma arginine only in male animals, with higher values in old animals relative to young animals (Fig S1A, p<0.05). There were no significant treatment effects on plasma phenylalanine levels (Fig S1A). Since insulin resistance is commonly linked to increased plasma BCAA and BCKA levels ^(9–11,43,44)^, we next looked at the effect of age and sex on insulin tolerance. There were no differences in basal glucose levels. At the initial critical time points of 0-15 min during the test, there no significant group differences, but old males and females had higher blood glucose levels at 120 min post insulin injection compared to their young counterparts (Fig 1C, p<0.05). This did not lead to a significant effect on area under the curve for glucose during the insulin tolerance test (ITT) between time points 0-120 min (Fig 1D). There was an age effect on body weight, as old animals exhibited higher body weight than their younger counterparts (Fig 1E, p<0.01). There was also an interaction effect, as old male mice had higher body weight than old female mice, an effect that was not seen in younger animals (Fig 1E, p<0.001).

### Skeletal muscle BCAA are reduced in old age while BCKA and relevant enzymes are unchanged

Skeletal muscle is a major contributor to whole-body BCAA oxidation ^(45)^, thus we looked at the BCAA/BCKA levels in skeletal muscle and the abundance/activity of some of the enzymes involved in BCAA metabolism. There was an interaction effect on total intracellular muscle BCAA levels, with old males having lower values compared to young counterparts (Fig 2A, p<0.01). There was significant interaction effect on leucine, and valine levels, with old male animals having lower values compared to young male animals (Fig 2A). In general, these trends are a reversal of what was observed for circulating BCAA (Fig 1A). There was no sex effect on total or any individual BCAA.

**Fig 2:**
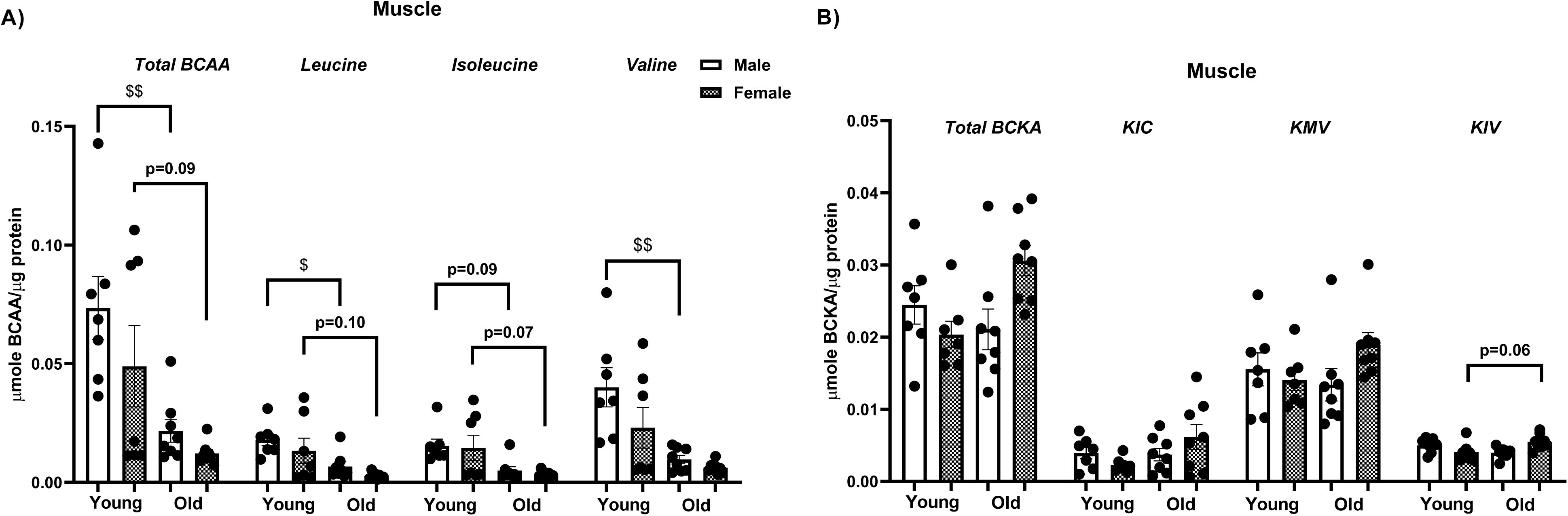
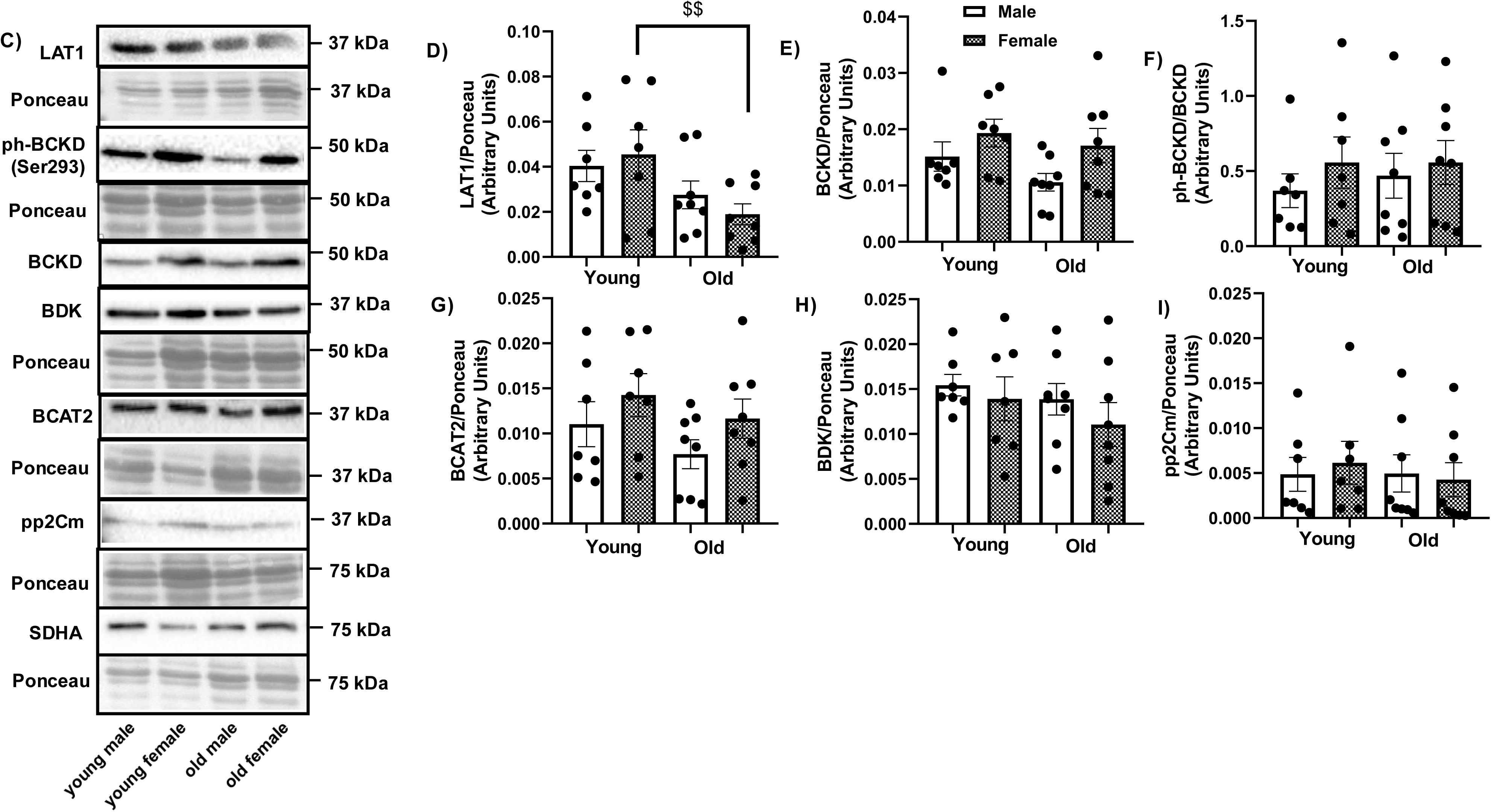
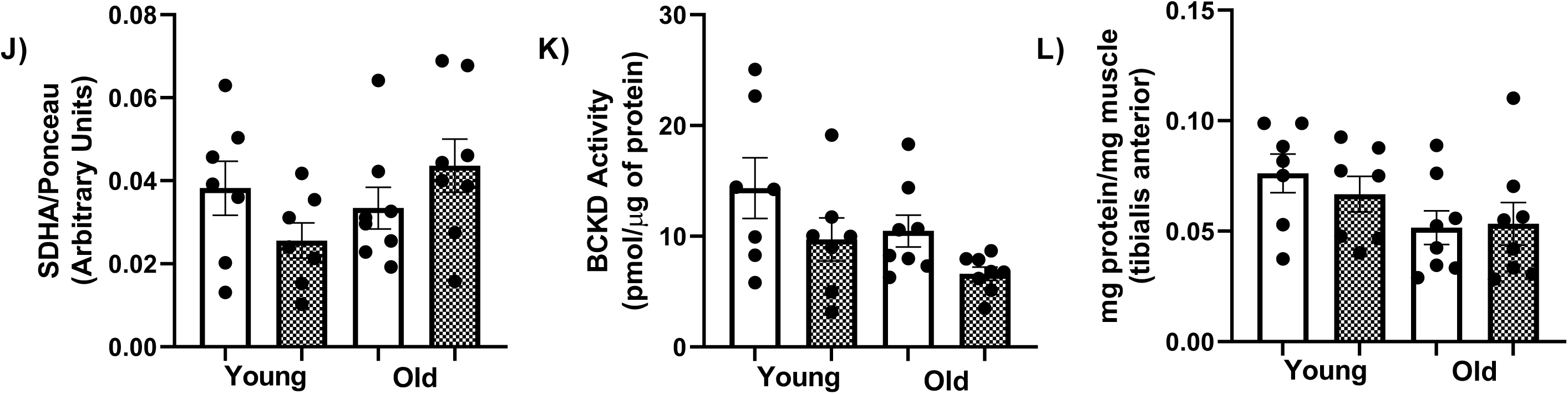
BCAA levels are reduced in muscle of older animals but with minimal changes in BCKA or BCAA catabolic enzymes. Muscle samples from young (n=7) and old (n=8) mice were analyzed by HPLC to measure intramuscular (tibialis anterior) total BCAA, leucine, isoleucine, valine (A), and total BCKA, KIV, KIC and KMV (B) concentrations. Proteins were immunoblotted against LAT1 (C-D), BCKD (C, E), phosphorylated (ph)-BCKD (Ser293) (C, F), BCAT2 (C, G), BDK (C, H), pp2Cm (C, I), and SDHA (C, J). BCKD activity assay was performed (K). Total weights of tibialis anterior (L), gastrocnemius (M), quadriceps (N), soleus (O), and EDL (P) muscles, expressed as per unit of per body weight. Protein concentration of the tibialis anterior normalized for its weight (Q).Data are means ± SEM. Effect of age: *** p<0.001, **** p<0.0001. Effect of age- sex interaction: $ p<0.05, $$ p<0.01, $$$ p<0.001.

There was no effect of age or sex on BCKA levels (Fig 2B) although old female tended (p=0.06) to have higher KIV compared to young female. There were no significant sex or age effect on muscle intracellular glutamate, serine, arginine, alanine, or phenylalanine levels (Fig S1B).

Changes in plasma and intramuscular BCAA levels could be attributed to changes in BCAA transporters like L-type amino acid transporter (LAT1). There was an age-sex interaction effect in that LAT1 levels were reduced in old females compared to young (Fig 2C-D, p<0.01), consistent with the general tendency for BCAA/Ile to be lower in old female (Fig 2A) There was no significant treatment effect for protein abundance for total BCKD, phosphorylated BCKD (Ser293), BCAT2, the BCKD kinase (BDK, a negative regulator of BCKD), and protein phosphatase 2Cm (pp2Cm, a positive regulator of BCKD) in skeletal muscle (Fig 2C-I). A lack of changes in BCAA catabolic enzymes could be due to differences in mitochondrial levels, as mitochondrial DNA volume, integrity, and functionality is reduced with aging ^(46)^, and these catabolic enzymes are mitochondrial proteins. Thus, we looked at succinate dehydrogenase (SDHA), a mitochondrial complex protein and a marker of mitochondrial abundance, but there was no significant effect of age or sex on SDHA protein levels in skeletal muscle (Fig 2C, J). There was no age or sex effect on BCKD activity (Fig 2K). Since reduced skeletal muscle mass is associated with reduced serum BCAA ^(47)^, we measured skeletal muscle weights. There was an age effect on weights of the tibialis anterior (Fig 2L, p<0.0001), gastrocnemius (Fig 2M, p<0.0001), quadricep (Fig 2N, p<0.0001), soleus (Fig 2O, p<0.001) muscles, as they were reduced in old animals. There was an interaction effect on tibialis anterior weight, as it was higher in old female compared to old male mice (Fig 2L, p<0.05). There was an interaction effect in the EDL, as old male mice had lower EDL weight than young male mice, (Fig 2P, p<0.001) and old female mice (Fig 2P, p<0.05). However, there was no significant treatment effect on muscle protein concentration normalized for the weight of the tibialis anterior, the muscle that was used for western blot analyses (Fig 2Q). Given the fact that muscle BCAA levels and LAT1 abundance are in general lower in old animals, the reduced muscle mass with age suggests that the effect of age on total BCAA and LAT1 is more pronounced than what is indicated by the muscle concentrations of these measures.

### In the liver, old animals have reduced pp2Cm independent of sex, while old female mice exhibit highest BCKA

Liver is an important contributor to BCAA catabolism, especially in the steps distal to BCKA ^(2)^. There was an age-sex interaction effect for reduced total BCAA and valine levels in old male animals compared to young male counterparts (Fig 3A, p<0.05). There were no significant group effects on liver leucine and isoleucine levels. There were age-sex interactions for liver BCKA levels in that total and individual BCKA values were the highest in old females, especially for KIV and KMV (Fig 3B).

**Fig 3:**
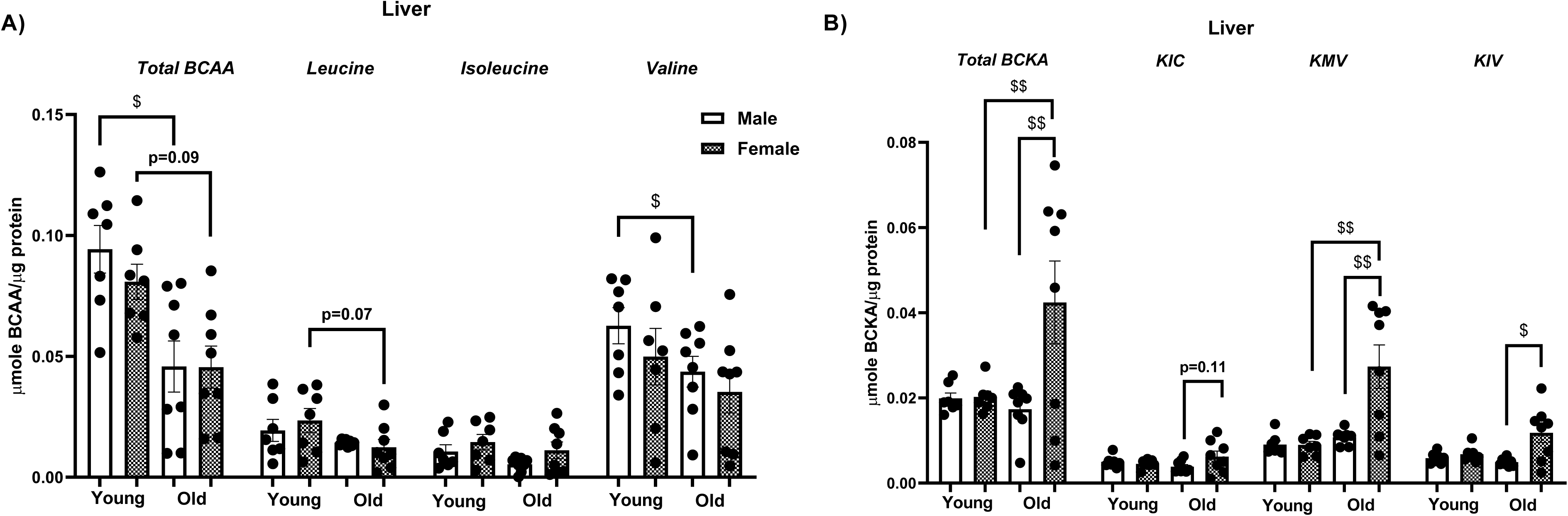
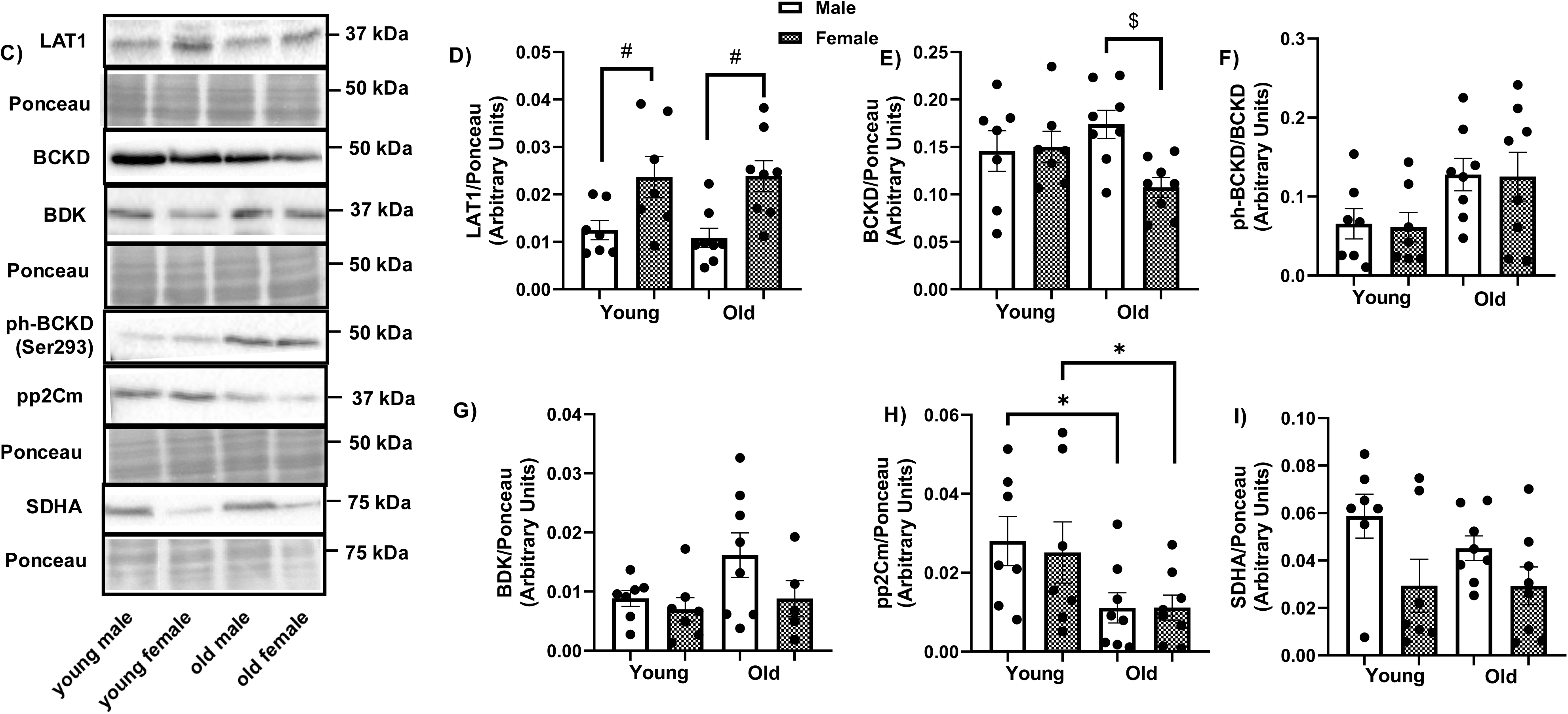
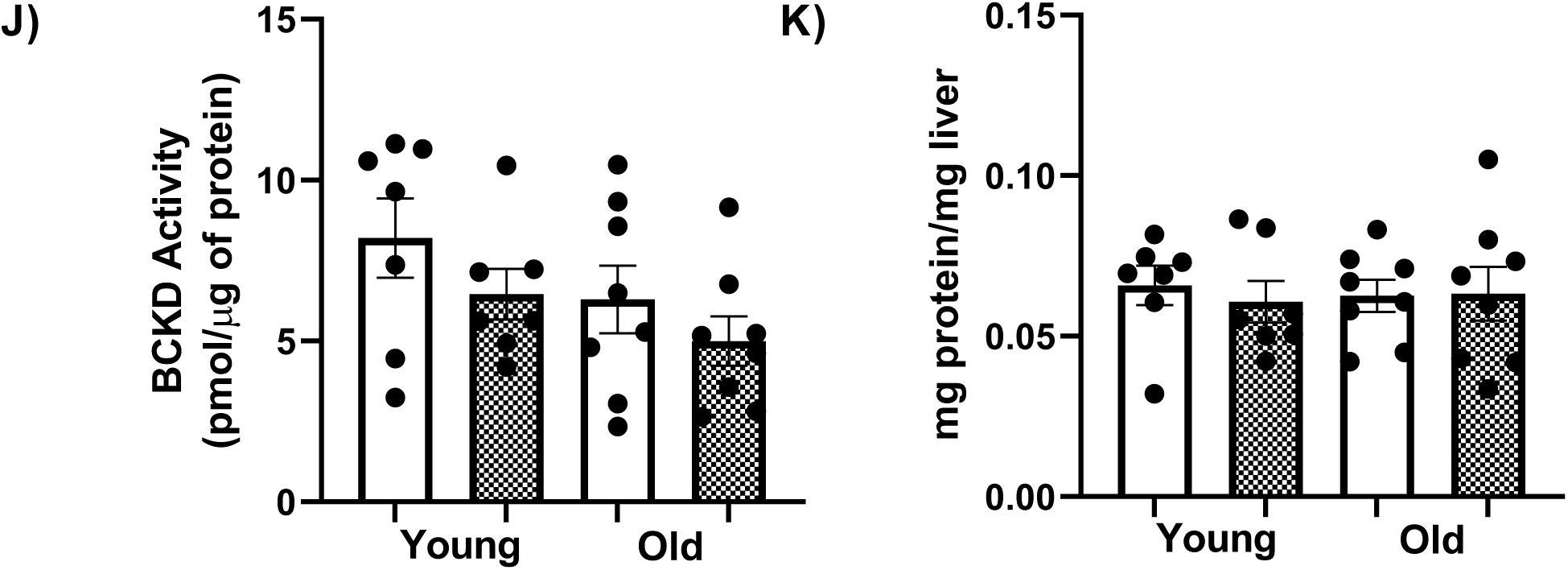
Reciprocal effects of age on liver BCAA and BCKA levels along with reduced pp2Cm abundance in old mice. Liver samples from young (n=7) and old (n=8) mice were analyzed by HPLC to measure liver total BCAA, leucine, isoleucine, valine (A), and total BCKA, KIV, KIC and KMV (B) concentrations. Proteins were immunoblotted against LAT1 (C-D), BCKD (C, E), ph-BCKD (Ser293) (C, F), BDK (C, G), pp2Cm (C, H), and SDHA (C, I). BCKD activity assay was performed (J). Weight of the liver per unit body weight (K). Protein concentration of the liver normalized for its weight (L). Data are means ± SEM. Effect of age: * p<0.05, *** p<0.001. Effect of sex: # p<0.05. Effect of age-sex interaction: $ p<0.05, $$ p<0.01.

There was no effect of sex or age on intracellular liver glutamate, serine or arginine levels (Fig S1C). There was an age-sex interaction effect on alanine levels as old male had reduced alanine compared to young male mice (Fig S1C, p<0.01). Phenylalanine levels were reduced in old animals irrespective of sex (Fig S1C, p<0.05).

There was an effect of sex on LAT1 protein abundance, as female mice had greater LAT1 abundance compared to male animals irrespective of age (Fig 3C-D, p<0.05). There was an interaction effect on total BCKD levels with the levels being reduced in old female as compared to male (Fig 3C, E, p<0.05). There was no effect of age or sex on the phosphorylation of BCKD (Ser293) nor on BDK (Fig 3C, F-G) p<0.01), but pp2Cm protein abundance was lower in old mice (Fig 3C, H, p<0.05). The reduced BCKD abundance along with reduced pp2Cm can at least in part explain the increased live BCKA levels in old mice. There was no sex or age effect on SDHA protein levels (Fig 3C, I) or BCKD activity (Fig 3J). There was an age effect on liver wet weight, as it was reduced in old compared to young mice (Fig 3K, p<0.05). There was also an interaction effect as old females had greater liver weight than old males (Fig 3K, p<0.05), but there was no significant effect on liver protein concentration (Fig 3L).

### In adipose tissue, BCKA are increased in old males along with a decrease in BCAT2 abundance in old female mice

Adipose tissue is another site of BCAA catabolism ^(48)^. There was no effect of age or sex on adipose tissue total BCAA, BCKA, or on any of the individual BCAA/BCKA (Fig 4A, B). However, there was an age-sex interaction effect on BCKA in that total BCKA, KIV and KMV in levels in old male mice were higher compared to young male mice (Fig 4B,). Also, in young but not the old animals, total BCKA and KIC levels were higher in female (Fig 4B).

**Fig 4:**
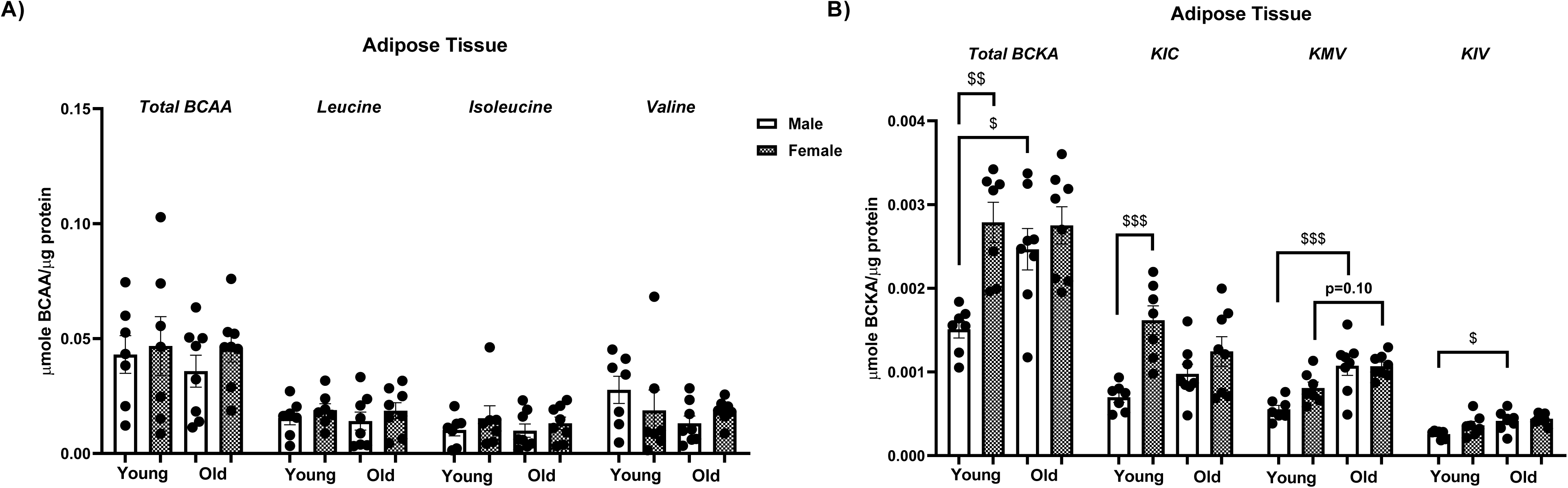
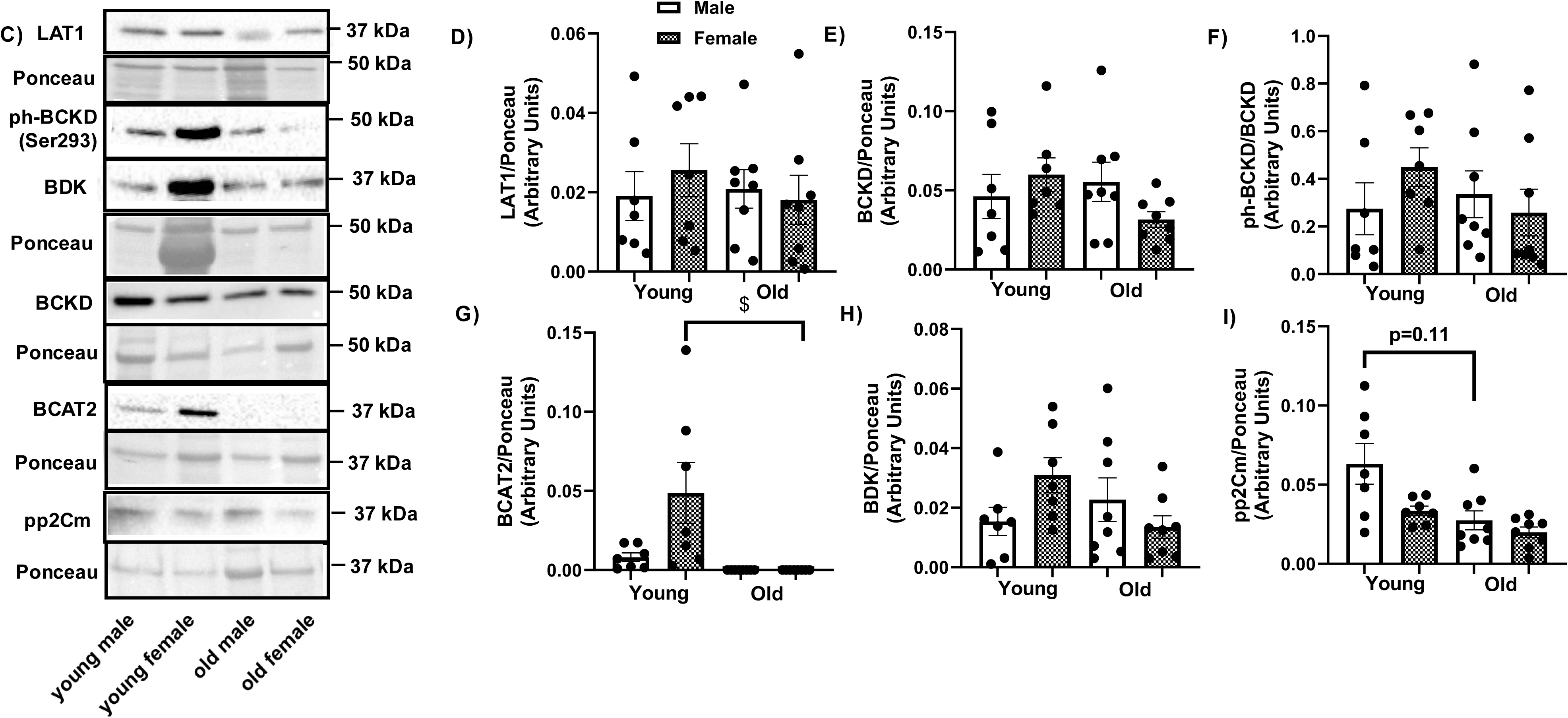
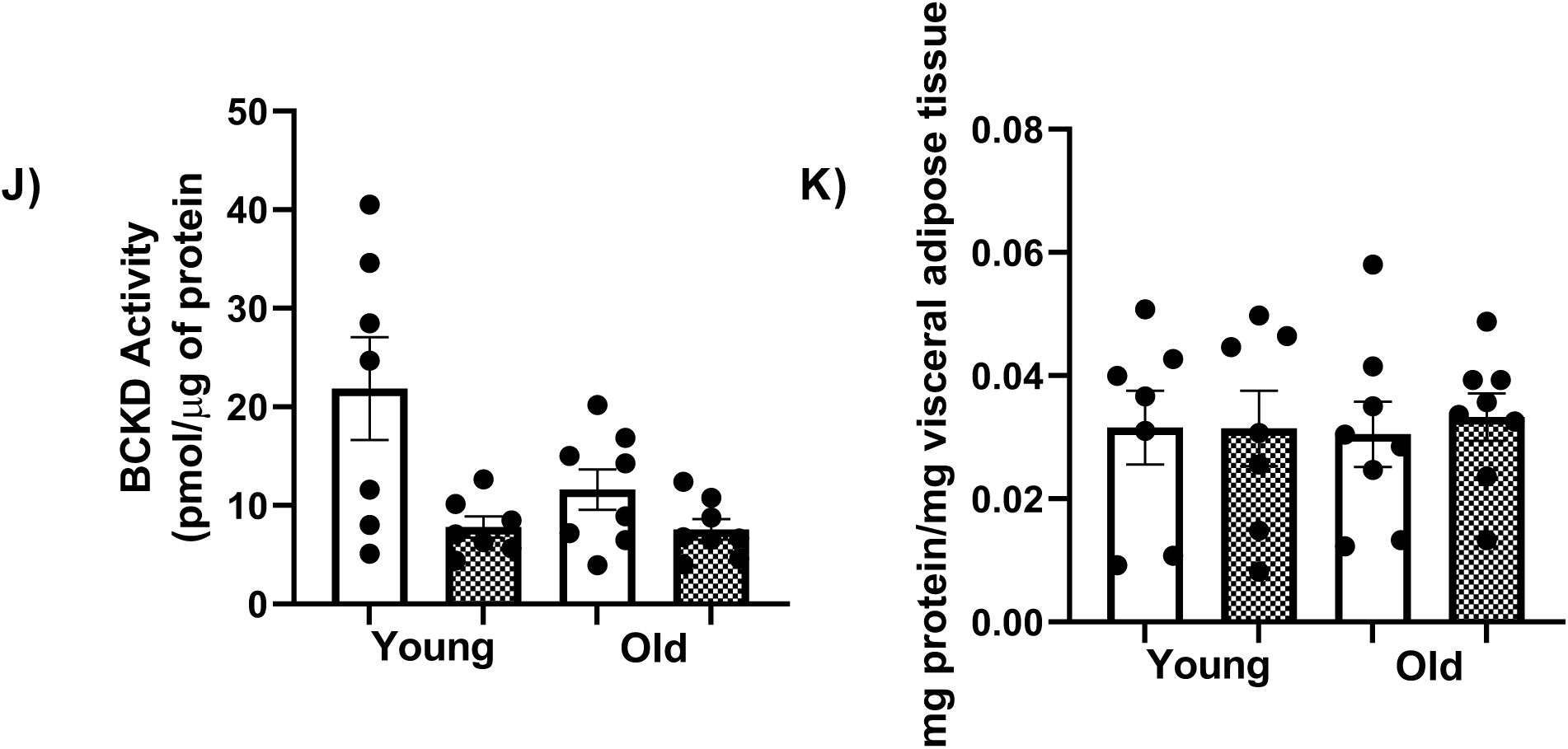
Age-sex interaction effects on BCKA, BCAT2 protein levels in mouse adipose tissue. Samples were as described in Fig 1-3. HPLC was performed to measure adipose tissue total BCAA, leucine, isoleucine, valine (A), and total BCKA, KIV, KIC and KMV (B) concentrations. Proteins were immunoblotted against LAT1 (C-D), BCKD (C, E), ph-BCKD (Ser293) (C, F), BCAT2 (C, G), BDK (C, H), pp2Cm (C, I). BCKD activity assay was performed (J). Total weight of visceral adipose tissue per unit body weight (K). Protein concentration in visceral adipose tissue normalized for its weight (L). Data are means ± SEM Effect of age: ** p<0.01, **** p<0.0001. Effect of age-sex interaction: $ p<0.05, $$ p<0.01, $$$ p<0.001.

There was no effect of age or sex on adipose tissue glutamate, serine, arginine, alanine, or phenylalanine levels (Fig S1D), nor on LAT1, total BCKD, phosphorylation of BCKD (Ser293), total BDK, pp2Cm, nor on BCKD activity (Fig 4C-F, H, I, J). However, in female, BCAT2 levels were lower in old compared to young female mice (Fig 4C, G, p<0.05). There was an age effect on visceral adipose tissue weight, with an increase in old animals (Fig 4K, p<0.01). There was also an interaction effect, as old female mice had greater visceral adipose tissue weight than old male mice (Fig 4K, p<0.01). However, there was no age or sex effect on tissue protein concentration normalized for the weight of the visceral adipose tissue (Fig 4L).

### BCAA are reduced in the heart in old age

Finally, we studied the heart as there is evidence for a link between impaired BCAA metabolism and heart failure and heart insulin resistance ^(49–51)^. In the heart, there was an effect of age on BCAA in that total BCAA, leucine, and valine levels were reduced in old animals (Fig 5A). For isoleucine, there was an age-sex interaction effect in that reduced level was observed only in old male relative to young male. There were also age-sex interactions on total BCAA and valine, as young females had reduced levels compared to young males (Fig 5A). There was no effect of age or sex on total or the individual BCKA levels (Fig 5B). There was no effect of age or sex on heart glutamate, serine, arginine, alanine and phenylalanine levels (Fig S1E).

**Fig 5:**
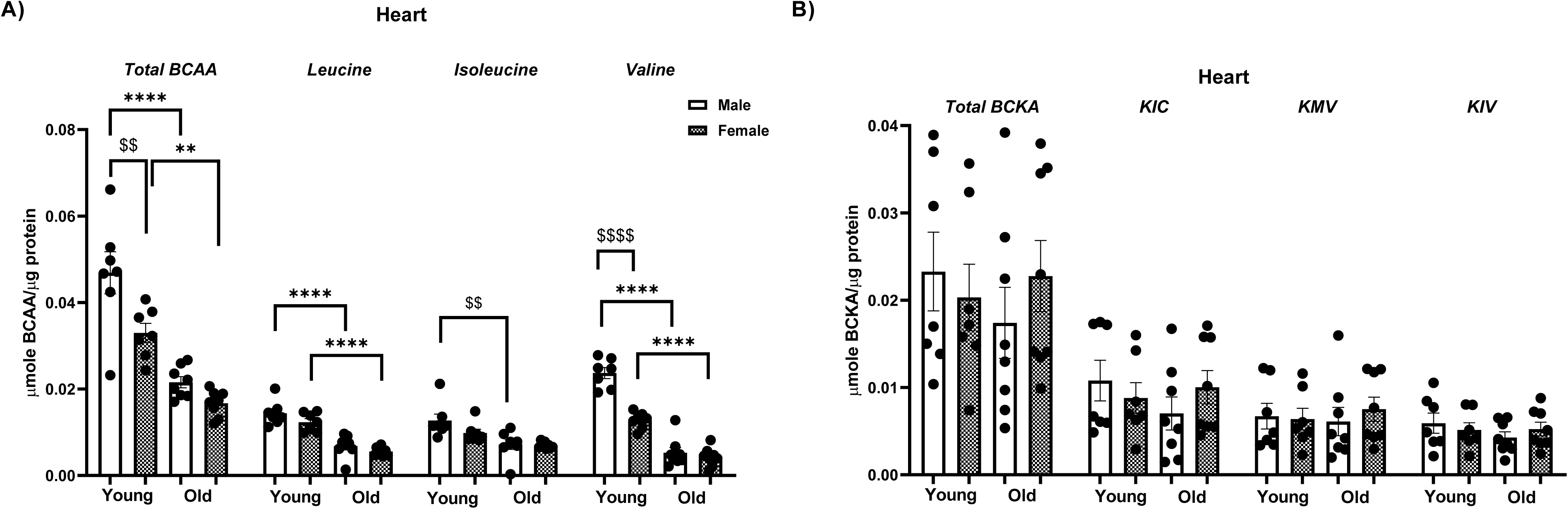
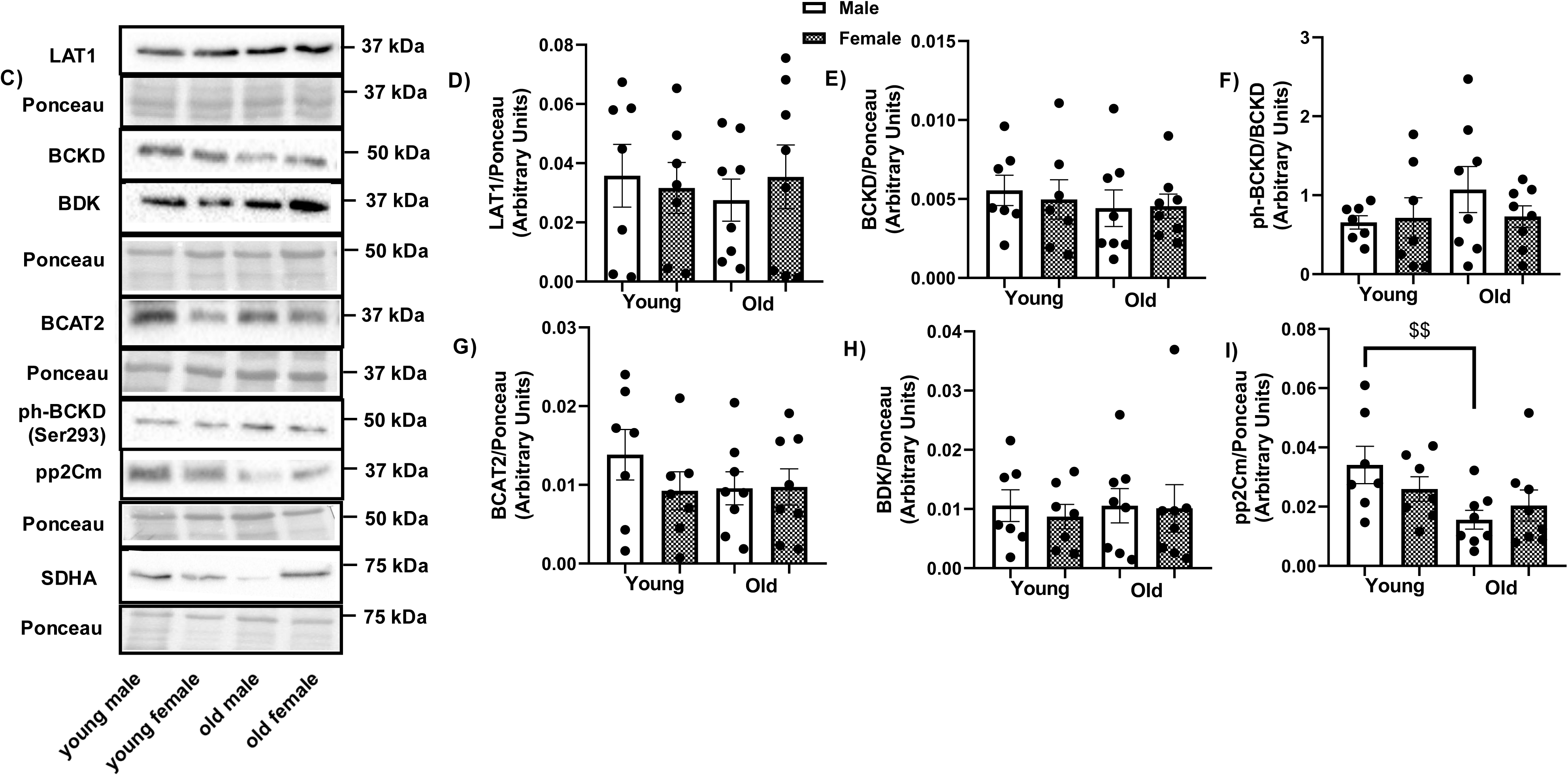
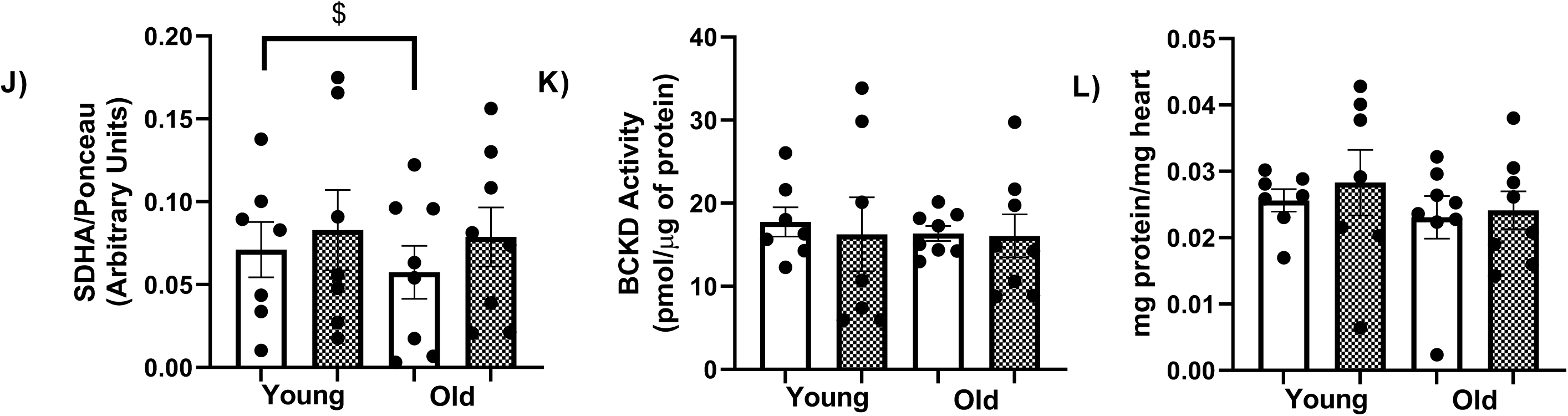
BCAA levels are reduced in the heart of old mice. Samples were as described in Fig 1-3. HPLC was performed to measure intracellular total BCAA, leucine, isoleucine, valine (A), and total BCKA, KIV, KIC and KMV (B) concentrations in the heart. Proteins were immunoblotted against LAT1 (C-D), BCKD (C, E), ph-BCKD (Ser293) (C, F), BCAT2 (C, G), BDK (C, H), pp2Cm (C, I), and SDHA (C, J). BCKD activity assay was performed (K). Total weight of the heart is expressed per unit of body weight (L). Protein concentration of the heart normalized for its weight (M). Data are means ± SEM. Effect of age: ** p<0.01, **** p<0.0001. Effect of age-sex interaction: $ p<0.05, $$ p<0.01, $$$ p<0.001, $$$$ p<0.0001.

In the heart, there was no effect of age or sex on LAT1, BCKD, ph-BCKD (Ser293), BCAT2, and BDK protein levels, nor on BCKD activity (Fig 5C-H, K). However, there was an interaction effect of age and sex on pp2Cm (Fig 5C, I, p<0.01) and SDHA (Fig 5C, J, p<0.05), as old males had reduced protein levels of these compared to young males. There was also an interaction effect on heart weight, as old female mice had reduced heart weight compared to young female (Fig 5L, p<0.001). Also, in the young groups, heart weight was higher in female (Fig 5L, p<0.05). There was no significant treatment effect on protein concentration normalized for the weight of the heart (Fig 5M).

### Plasma BCAA and BCKA levels are poorly correlated with basal plasma glucose levels

There were tendencies for weak positive correlations between basal plasma glucose levels and plasma total and individual BCAA. The associations with the BCKA were very weak (Fig 6A-H).

**Fig 6:**
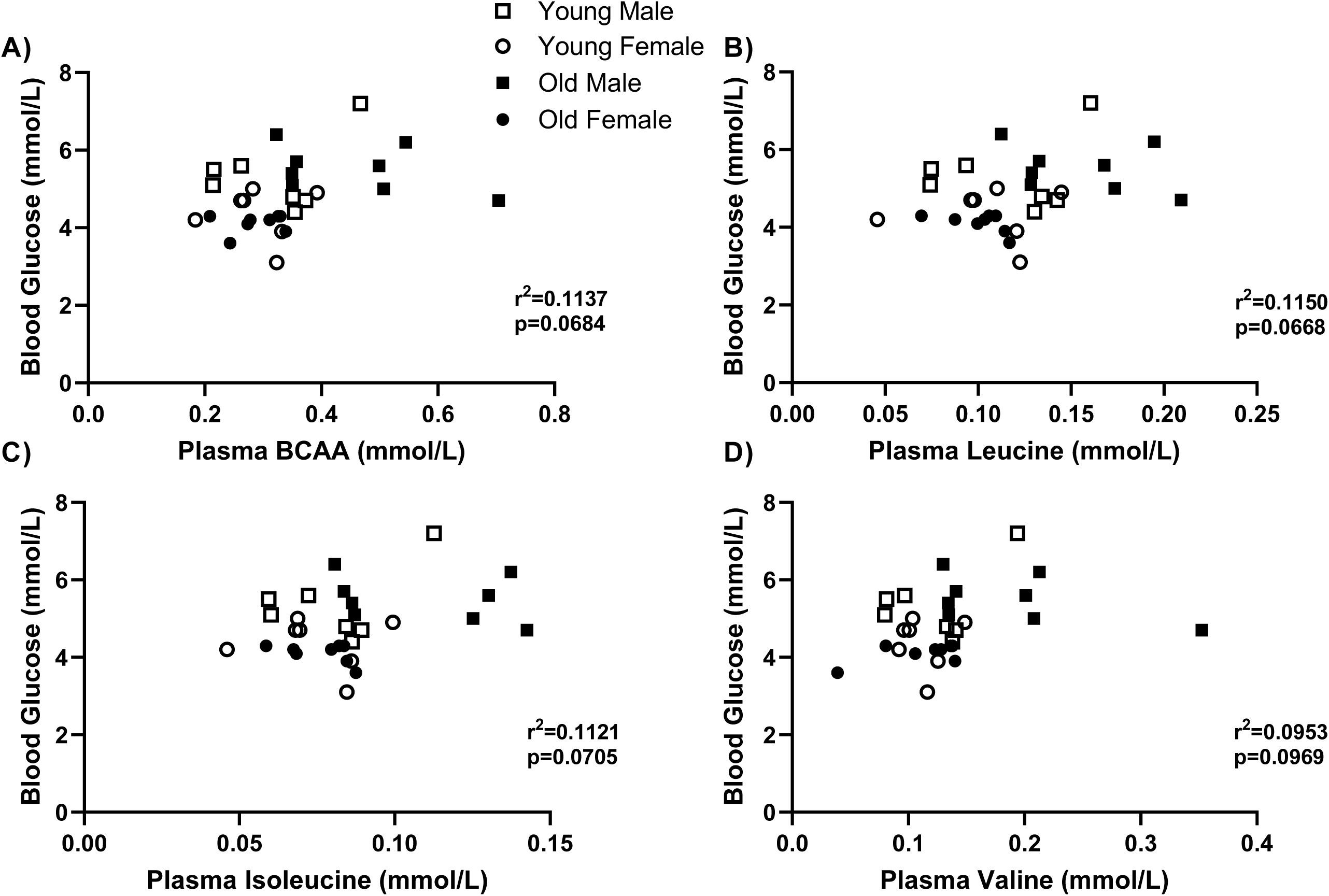
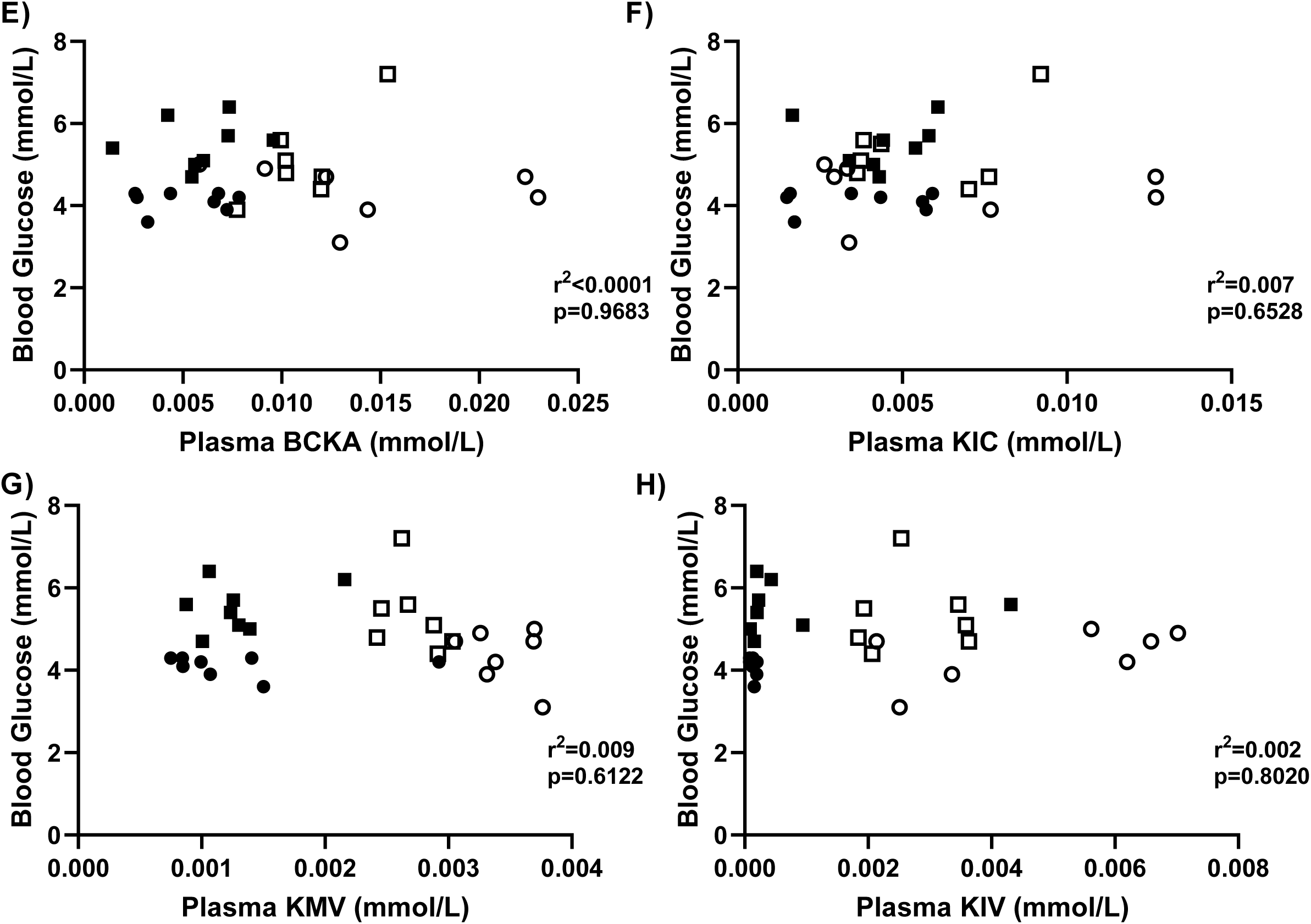
Plasma BCAA and BCKA and basal plasma glucose levels are poorly correlated. Correlations between basal plasma glucose levels and plasma total BCAA (A), leucine (B), isoleucine (C), valine (D), total BCKA (E), KIC (F), KMV (G) and KMV (H). Data were analyzed using multiple linear regression. These charts are drawn from the data in Fig 1.

## Discussion

Elevated circulating BCAA and defects in BCAA metabolism are linked to a plethora of diseases/conditions like insulin resistance, liver cirrhosis, chronic kidney failure and maple syrup urine disease ^(3–11)^, and aging is a major risk factor in many of these diseases ^(52–55)^. Thus, it would be expected that circulating (and tissue) levels of BCAA would increase with age, and that aging would be associated with impaired BCAA catabolism. Because this relationship could be modified by sex, here we studied the effects of age and sex on circulating BCAA and BCKA, and on tissue levels and metabolism (oxidation) of these amino acids. We demonstrated that alone, age or sex has limited effects on many of the measures. For example, there was reduced circulating BCKA in old animals along with reduced muscle and liver total BCAA levels in old animals. On the other hand, muscle LAT1 abundance was greater in female irrespective of age. Notwithstanding these effects of age or sex, many of the other changes that we observed are a result of the interactions of age and sex. We also note that in general, circulating levels of the BCAA/BCKA do not predict tissue levels of these metabolites. Changes in BCAA/BCKA content could not be attributed to significant changes in BCKD activity in any of the tissues. Finally, circulating levels of BCAA and BCKA correlated poorly with basal blood glucose levels. These data suggests that the capacity to catabolize BCAA is preserved in aging in healthy animals and is likely only dysregulated in disease states.

BCAA levels in the body are regulated by a balance between their inputs (such as dietary intake, synthesis by gut microbes, and protein breakdown) and their disappearance (including incorporation into proteins, excretion, and metabolic breakdown) ^(1,56)^. From previous studies, there does not appear to be any universality with regards to the effect of age on circulating BCAA. Circulating BCAA levels are lower in men as they age ^(24,25)^. Elderly individuals (average 80 years of age) have reduced plasma leucine and isoleucine levels compared to their young counterparts (average 29 years of age) ^(57)^. Conversely in another study, age did not impact total serum amino acid levels or serum BCAA levels, but there was greater serum BCAA levels in males compared to females independent of age ^(58)^. Our observation of limited effect of age on circulating BCAA is in general consistent with this study. However, in one study serum BCAA are higher in an old male population compared to old females, which correlated with reduced energy and protein intake in the female population ^(59)^. Studies in mice showed higher ^(28)^, a trend for higher ^(38)^ or no difference ^(39)^ in plasma BCAA in aging. Generally, in old age, protein intake is reduced ^(60)^, which could explain the reduction in BCAA in old men ^(24,25,61)^. The old male mice from this study consumed more food than old female mice (∼18%), but when corrected for body weight the difference was negligible. Young mice did consume more food per day than old mice (∼20%), which could also influence intramuscular BCAA content, but this was not consistent with changes in plasma BCAA in females. One study showed no link between protein intake and serum BCAA concentrations in healthy males, but an inverse relationship in healthy females ^(62)^. Interestingly, they also show that plasma BCAA concentration is inversely correlated with losses in skeletal mass index in healthy males ^(62)^. Here we show that plasma BCAA are increased in old male mice consistent with reduced tibialis anterior (Fig 2L), gastrocnemius (Fig 2M), quadricep (Fig 2N), soleus (Fig 2O and EDL (Fig 2P) muscle weight in old males compared to young male counterparts.

The differences observed in circulating and tissue BCAA levels would be due to aggregated effects of the prandial state of the animals, adaptation to habitual diets, and integrative metabolism in tissues. Because samples were collected in the post absorptive state and the animals were fed the same diets (albeit for different length of time because of the differences in age), the observed differences would be due more to integrative interorgan metabolism of these amino acids rather than being a reflection of acute food intake. This metabolism would reflect incorporation of the amino acids into tissue proteins, release from tissue proteins (proteolysis), transport into and out of the tissue, and metabolism (transamination, oxidation, use to make other compounds).

Changes in tissue BCAA/BCKA levels can be affected by tissue wasting and protein turnover. Decline in muscle mass is associated with reductions in serum BCAA ^(47)^. Consistent with this, reductions in muscle BCAA in old animals occurred in parallel with a reduction in tibialis anterior weight (Fig 2L). In muscle wasting conditions (like trauma and sepsis), BCKA are converted back to BCAA ^(63)^, which can serve to prevent further fall in muscle BCAA. Changes in basal protein degradation/synthesis could affect BCAA/BCKA levels in muscle. There are inconsistencies on the effect of age on measures of muscle protein synthesis and degradation, with some studies reporting no change in basal protein synthesis between young and elderly male individuals ^(64–66)^ while others reported reduced basal muscle protein synthesis in old animals compared to their young counterparts ^(67,68)^. Regarding proteolysis, one study reported increases in skeletal muscle mRNA content of drivers of proteolysis. including forkhead box protein O3a (FoxO3a) and muscle ring-finger protein-1 (MuRF-1) in old women compared to young women ^(69)^ while another study reported downregulation of mRNA content of MuRF-1 and atrogin-1 in skeletal muscle of old female rats compared to younger counterparts ^(70)^. In their study, Welle et al. found no effect of age on E3 ligases mRNA content in male rats ^(71)^. These ambiguities on the effect of age on indicators of muscle protein turnover make it challenging to speculate on the interactions between muscle protein turnover and BCAA concentrations/metabolism during aging.

Age-related changes in renal extraction may affect circulating BCAA. Glomerular filtration rate (GFR) is reduced in aging (80). This corresponded with reduced reabsorption and enhanced excretion of leucine but not valine and isoleucine. However, this did not correspond with any changes in serum leucine levels in aging. This study did not measure if other mechanisms (BCAA metabolism, protein synthesis/breakdown) had any compensatory effects (80). Thus, it is unlikely that possible age-related reduced GFR may explain our data because we observed increased circulating BCAA levels in old male mice.

BCAA biosynthesis by gut bacteria is increased in insulin resistance and obesity as reviewed by Gojda et al (81). Despite no change in our ITT data, adipose tissue weight was increased in old animals, which perhaps could explain the increase in plasma BCAA in old male animals. However, this is not consistent with a lack of change in plasma BCAA levels in female mice despite increases in adipose tissue weight in old female mice. Additionally, estrogen inhibits BCAA oxidation (29,30), however, contrary to increased circulating BCAA levels that one might expect in female animals, old female animals had reduced levels.

Inflammation is increased with aging ^(72)^. Therefore, altered metabolism of the BCAA and other amino acids in aging may be related to this condition. Glutamate-glutamine metabolism is linked to BCAA metabolism. We only observe a trend (P=0.09) for reduced muscle glutamate in old animals, especially in old female relative to young female. Depletion of glutamate could be due to their conversion to glutamine, as glutamine inhibits inflammation in muscle ^(73)^ and is important for the functioning of the immune system ^(74)^. Because muscle samples were collected in the post absorptive state, another reason for reduced glutamate level might be due to its use as a substrate for the generation of α-ketoglutarate, an anaplerotic substrate of the TCA circle.

As mentioned before, altered tissue metabolism would contribute to the differences in tissue circulating levels of BCAA/BCKA. Metabolism of the BCAA is related to the activities of the relevant enzymes, which would be affected by the abundance of the enzymes and relevant posttranslational modifications. In this study, except for changes in LAT1 in muscle and liver, there were no consistent significant effects of age or sex on these enzymes. This could be due to altered mitochondrial content, as these BCAA catabolic enzymes are present in the mitochondria, and mitochondrial DNA volume, integrity and function are reduced with aging ^(46)^. There are reduced oxidative phosphorylation (OXPHOS) activity, elevated oxidative damage, reduced mitochondrial quality control, decreased activity of metabolic enzymes, as well as alterations in mitochondrial morphology, dynamics and biogenesis in aging ^(75,76)^. In our study, only in the heart was the mitochondrial abundance (as determined by SDHA level) reduced in old male relative to young male mice. There was no effect of age, sex, nor the interaction of the two on mitochondrial content in muscle or any of the other tissues. This could be due to the muscle analyzed, as in the vastus lateralis muscle, old animals had reduced complex III, but not complex I, II, IV or V protein abundance compared to young counterparts ^(77)^. Conversely, one study shows that in the gastrocnemius and EDL the abundance of mitochondrial complex proteins was increased in old age, while in the soleus, there was a reduction in complex I and IV protein abundance and no change in the adductor longus ^(78)^. Additionally, another study shows a transient dip in mitochondrial density in old age (24-month rats), but this returns to normal in even older age (35-month rats) ^(79)^. Therefore, it is possible that the effect of age on mitochondrial content might depend on the model of aging, the type of muscle and the specific mitochondrial proteins that are analyzed. This makes it difficult to attribute any changes in BCAA catabolic enzyme activity and abundance during aging to changes in mitochondria content. Future studies measuring mitochondrial respiration, ROS generation, and mitochondrial DNA content may shed more light on the link between mitochondria and changes in these BCAA enzyme levels and BCAA metabolism.

The effects of sex on many of the parameters measured were seen only in the context of age-sex interactions. For example, circulating total BCAA as well as leucine and valine levels were elevated in old male relative to old female: this sex effect was not seen in young animals. Related, while there was a clear effect of sex on liver LAT1, the effect of age on muscle LAT1 was particularly obvious when comparing young female to old female. The sex hormone estrogen inhibits the BCKD complex by activating BDK, thus decreasing BCAA catabolism ^(29,30)^. However, this did not translate to changes in BCAA catabolic enzyme abundance or activity. In old age, there should be minimal sex differences on estrogen level. There was only a sex effect on protein abundance of LAT1 abundance in the liver, as it was higher in females compared to males. We also saw age-sex interactions on muscle LAT1, with LAT1 abundance being reduced in old female relative to young female, something that was not observed for male. From the effect of estrogen on BDK, the overall expectation would be for reduced BCKD activity in female and attendant increases in tissue and circulating BCAA/BCKA. This was the case for only liver total BCKA, KIV and KMV. This moderate effect is likely due to the fact that BCKD activity and BCAA oxidation can be regulated at the levels of other enzymes in the BCAA catabolic pathway.

In a study that examined sex effects on BCAA levels in humans ^(80)^, men (median age 57) had higher plasma BCAA levels than women (median age 60), which is consistent with our data. These differences could not be explained by differences in dietary protein intake ^(80)^. Interestingly, men have greater leucine oxidation compared to women following endurance exercise ^(32)^. Another study showed that adolescent women with T2D had greater BCAA/BCKA excretion than adolescent males. This sex difference was not seen in obese non-diabetic adolescents ^(81)^, collectively suggesting context dependency of the effect of sex on BCAA oxidation. Two studies reported higher muscle protein synthesis in old women compared to BMI and age matched men, despite women having less muscle mass ^(82,83)^, an observation that appears to hold true only later in life, but not in middle aged men and women ^(84)^. If the same is true in old mice, better utilization of BCAA for protein synthesis could explain reduced circulating BCAA levels observed in old female mice.

Reduced BCAT2 protein levels were observed in adipose tissue of old male mice compared to young male mice, which correlated with increased plasma BCAA concentrations in old male mice ^(28)^ just like in our study. Although we did not see group effect on adipose tissue BCAT2 level, BCAT2 bands were only barely visible in old mice, suggesting the possibility of a type II error.

Increased circulating BCAA and BCKA are linked to insulin resistant states ^(9–11,17–21)^. In older populations, insulin sensitivity is reduced ^(26,85,86)^. We observed no group effect on blood glucose level at the relevant time points 0-15 min of the ITT. While there was a group effect on blood glucose at 120 min, this was likely due to the activation of the counter regulatory system. The lack of significant differences in our ITT data is consistent with the lack of changes in BCAA catabolic enzyme abundance as BCAA catabolic protein abundance is downregulated in insulin resistance in muscle ^(87)^, liver ^(88)^ and adipose tissue ^(11)^.

What do the age/sex effects on BCAA levels and metabolism imply? Increased plasma BCAA in old male mice might serve a compensatory mechanism to increase life span, as endogenous BCAA accumulation has a protective effect on leukocyte telomere length and weakness in an east Asian population (56). Also, BCAA supplementation increased life span of male mice (16 months) by 12% (57). However, BCAA restriction increased lifespan in Drosophila (58). This could be by reducing ribosomal protein S6 kinase (S6K1) phosphorylation, as BCAA are potent activators of S6K1, and downregulation of S6K1 prolongs lifespan of mammals (59). Despite increased plasma concentrations in old male mice, the reduced intramuscular BCAA content in old male mice compared to young males should prevent S6K1 activation, leading to a prolonged lifespan.

A limitation of this study is that the animals were studied in the postabsorptive state in order to capture the effects of chronic regulation of BCAA metabolism in tissues. Studying the animals in the fed state might provide additional insights into how the interactions of age and sex can shape BCAA metabolism and associated age-related metabolic abnormalities. Furthermore, although 18-month mice are considered “old” ^(89)^, inclusion of older animals (≥ 24 months) can help to establish whether there is a gradient in terms of the link between BCAA metabolism and aging but that was not captured in the current study. This is because abnormalities such as insulin resistance ^(86)^, worsening muscle atrophy ^(90)^, and reduced mitochondrial content ^(91)^ are likely to be seen in very old animals so it might be possible to establish whether these are linked to impaired BCAA metabolism. Lastly, while we observed statistically significance differences in some key measures (including BCAA levels in blood and tissues), the p values for some of the measures were borderline values required for statistical significance. Therefore, the possibility of type II errors exists for some of the measures. Future studies with larger sample size will help to address this.

In conclusion, we showed that rather than isolated effects of age or sex, these two factors interact in regulating local and systemic BCAA metabolism. Whereas there are age and/sex effects on tissue BCAA levels, these are not reflected in the circulating levels of these amino acids. The changes in BCAA and or BCAA cannot be predicted from the changes to the abundance/regulation of enzymes involved in BCAA catabolism. All these point to the complex interactions between the diverse relevant enzyme in shaping tissue BCAA metabolism. The net effects of such interactions will further be modulated by integrated metabolism across the different tissues as well as by factors such as the fed state of the individual, the degree of aging, metabolic state (for example, exercise) and health status.

## Acknowledgments

The authors thank the Muscle Health Research Centre at York University for use of the HPLC and Dr. Paluzzi’s lab for the use of their imaging systems. They also thank the Faculty of Health, York University for additional support.

## Authorship

GM and OAJA conceived and designed the experiments. GM and SM performed the experiments. GM drafted the initial version of the manuscript. OAJA reviewed and edited the manuscript. All authors approved the final version of the manuscript.

## Supporting Information

**Fig S1: Effects of age and sex on plasma and tissue glutamate, serine, arginine, alanine, and phenylalanine levels.**

Male and female young (4-month) (n=7) and old (18-months) (n=8) mice were sacrificed. HPLC was then performed to measure glutamate, serine, arginine, alanine, and phenylalanine in the plasma (A), muscle (B), liver (C), visceral adipose tissue (D), and heart (E). Data are means ± SEM. Effect of age: * p<0.05, ** p<0.01, **** p<0.0001. Effect of age-sex interaction: $ p<0.05, $$ p<0.01, $$$ p<0.001.

**Fig S2: Insulin tolerance test in young and old mice.**

**Fig S3: Skeletal muscle weights in young and old mice**

## References

1. Lynch CJ, Adams SH. Branched-chain amino acids in metabolic signalling and insulin resistance. Nat Rev Endocrinol. 2014;10(12):723–36.

2. Harper a E, Miller RH, Block KP. Branched-Chain Amino Acid Metabolism. Metabolism. 1984;30(6):409–54.

3. Indo Y, Kitano A, Endo F, Akaboshi I, Matsuda I. Altered kinetic properties of the branched-chain α-keto acid dehydrogenase complex due to mutation of the β-subunit of the branched-chain α-keto acid decarboxylase (E1) component in lymphoblastoid cells derived from patients with maple syrup urine disease. J Clin Invest. 1987;80(1):63–70.

4. Cano NJM, Fouque D, Leverve XM. Application of branched-chain amino acids in human pathological states: Renal failure. J Nutr. 2006;136(1):299–307.

5. Holecek M, Kandar R, Sispera L, Kovarik M. Acute hyperammonemia activates branched-chain amino acid catabolism and decreases their extracellular concentrations: Different sensitivity of red and white muscle. Amino Acids. 2011;40(2):575–84.

6. Tajiri K, Shimizu Y. Branched-chain amino acids in liver diseases. Vol. 3, World Journal of Gastroenterology. 2018. p. 47.

7. Blackburn PR, Gass JM, Pinto e Vairo F, Farnham KM, Atwal HK, Macklin S, et al. Maple syrup urine disease: Mechanisms and management. Appl Clin Genet. 2017;10:57– 66.

8. Xu J, Jakher Y, Ahrens-Nicklas RC. Brain branched-chain amino acids in maple syrup urine disease: Implications for neurological disorders. Int J Mol Sci. 2020;21(20):1–18.

9. Andersson-Hall U, Gustavsson C, Pedersen A, Malmodin D, Joelsson L, Holmäng A. Higher concentrations of BCAAs and 3-HIB are associated with insulin resistance in the transition from gestational diabetes to type 2 diabetes. J Diabetes Res. 2018;2018:1–12.

10. Giesbertz P, Padberg I, Rein D, Ecker J, Höfle AS, Spanier B, et al. Metabolite profiling in plasma and tissues of ob/ob and db/db mice identifies novel markers of obesity and type 2 diabetes. Diabetologia. 2015;58(9):2133–43.

11. Lackey DE, Lynch CJ, Olson KC, Mostaedi R, Ali M, Smith WH, et al. Regulation of adipose branched-chain amino acid catabolism enzyme expression and cross-adipose amino acid flux in human obesity. Am J Physiol - Endocrinol Metab. 2013;304(11):1175– 87.

12. Hara Y, May RC, Kelly RA, Mitch WE. Acidosis, not azotemia, stimulates branched-chain, amino acid catabolism in uremic rats. Kidney Int. 1987;32(6):808–14.

13. Mitch WE, Price SR, May RC, Jurkovitz C, England BK. Metabolic Consequences of Uremia: Extending the Concept of Adaptive Responses to Protein Metabolism. Am J Kidney Dis. 1994 Feb;23(2):224–8.

14. Shimomura Y, Honda T, Shiraki M, Murakami T, Sato J, Kobayashi H, et al. Branched-chain amino acid catabolism in exercise and liver disease. In: Journal of Nutrition. American Institute of Nutrition; 2006. p. S250–3.

15. Hernández-Alvarez MI, Díaz-Ramos A, Berdasco M, Cobb J, Planet E, Cooper D, et al. Early-onset and classical forms of type 2 diabetes show impaired expression of genes involved in muscle branched-chain amino acids metabolism. Sci Rep. 2017;7(1):1–12.

16. Lian K, Du C, Liu Y, Zhu D, Yan W, Zhang H, et al. Impaired adiponectin signaling contributes to disturbed catabolism of branched-chain amino acids in diabetic mice. Diabetes. 2015;64(1):49–59.

17. Mann G, Adegoke OAJ. Effects of ketoisocaproic acid and inflammation on glucose transport in muscle cells. Physiol Rep. 2021;9(1):e14673.

18. Moghei M, Tavajohi-Fini P, Beatty B, Adegoke OAJ. Ketoisocaproic acid, a metabolite of leucine, suppresses insulin-stimulated glucose transport in skeletal muscle cells in a BCAT2-dependent manner. Am J Physiol - Cell Physiol. 2016;311(3):C518–27.

19. Macotela Y, Emanuelli B, Bång AM, Espinoza DO, Boucher J, Beebe K, et al. Dietary leucine - an environmental modifier of insulin resistance acting on multiple levels of metabolism. PLoS One. 2011;6(6):e21187.

20. Tremblay F, Brûlé S, Sung HU, Li Y, Masuda K, Roden M, et al. Identification of IRS-1 Ser-1101 as a target of S6K1 in nutrient- and obesity-induced insulin resistance. Proc Natl Acad Sci U S A. 2007;104(35):14056–61.

21. Tremblay F, Marette A. Amino acid and insulin signaling via the mTOR/p70 S6 kinase pathway. A negative feedback mechanism leading to insulin resistance in skeletal muscle cells. J Biol Chem. 2001;276(41):38052–60.

22. Lerin C, Goldfine AB, Boes T, Liu M, Kasif S, Dreyfuss JM, et al. Defects in muscle branched-chain amino acid oxidation contribute to impaired lipid metabolism. Mol Metab. 2016;5(10):926–36.

23. White PJ, McGarrah RW, Grimsrud PA, Tso SC, Yang WH, Haldeman JM, et al. The BCKDH Kinase and Phosphatase Integrate BCAA and Lipid Metabolism via Regulation of ATP-Citrate Lyase. Cell Metab [Internet]. 2018;27(6):1281–1293.e7. Available from: 10.1016/j.cmet.2018.04.015

24. Ribeiro R V., Solon-Biet SM, Pulpitel T, Senior AM, Cogger VC, Clark X, et al. Of older mice and men: Branched-chain amino acids and body composition. Nutrients. 2019;11(8).

25. Sun L, Hu C, Yang R, Lv Y, Yuan H, Liang Q, et al. Association of circulating branched-chain amino acids with cardiometabolic traits differs between adults and the oldest-old. Oncotarget. 2017;8(51):88882–93.

26. Shou J, Chen PJ, Xiao WH. Mechanism of increased risk of insulin resistance in aging skeletal muscle. Diabetol Metab Syndr [Internet]. 2020;12(1):1–10. Available from: 10.1186/s13098-020-0523-x

27. Krok-Schoen JL, Archdeacon Price A, Luo M, Kelly OJ, Taylor CA. Low Dietary Protein Intakes and Associated Dietary Patterns and Functional Limitations in an Aging Population: A NHANES Analysis. J Nutr Heal Aging. 2019;23(4):338–47.

28. Han HS, Ahn E, Park ES, Huh T, Choi S, Kwon Y, et al. Impaired BCAA catabolism in adipose tissues promotes age-associated metabolic derangement. Nat Aging. 2023;3:982–1000.

29. Obayashi M, Shimomura Y, Nakai N, Jeoung NH, Nagasaki M, Murakami T, et al. Estrogen controls branched-chain amino acid catabolism in female rats. J Nutr. 2004;134(10):2628–33.

30. Kobayashi R, Shimomura Y, Murakami T, Nakai N, Otsuka M, Arakawa N, et al. Hepatic branched-chain α-keto acid dehydrogenase complex in female rats: Activation by exercise and starvation. J Nutr Sci Vitaminol (Tokyo). 1999;45(3):303–9.

31. Costanzo M, Caterino M, Sotgiu G, Ruoppolo M, Franconi F, Campesi I. Sex differences in the human metabolome. Biol Sex Differ. 2022;13(30):1–18.

32. Mittendorfer B, Horowitz JF, Klein S. Effect of gender on lipid kinetics during endurance exercise of moderate intensity in untrained subjects. Am J Physiol - Endocrinol Metab. 2002;283:E59–65.

33. Burke PJ, Karp JE, Vaughan WP. Chemotherapy of leukemia in mice, rats, and humans relating time of humoral stimulation, tumor growth, and clinical response. J Natl Cancer Inst. 1981;67(3):529–38.

34. Egorin MJ, Sentz DL, Rosen DM, Ballesteros MF, Kearns CM, Callery PS, et al. Plasma pharmacokinetics, bioavailability, and tissue distribution in CD2F1 mice of halomon, an antitumor halogenated monoterpene isolated from the red algae Portieria hornemannii. Cancer Chemother Pharmacol. 1996;39:51–60.

35. Farhang-Sardroodi S, La Croix MA, Wilkie KP. Chemotherapy-induced cachexia and model-informed dosing to preserve lean mass in cancer treatment. PLoS Comput Biol. 2022;18(3):e100950.

36. Essex AL, Pin F, Huot JR, Bonewald LF, Plotkin LI, Bonetto A. Bisphosphonate Treatment Ameliorates Chemotherapy-Induced Bone and Muscle Abnormalities in Young Mice. Front Endocrinol (Lausanne). 2019;10:809.

37. Eiseman JL, Eddington ND, Leslie J, MacAuley C, Sentz DL, Zuhowski M, et al. Plasma pharmacokinetics and tissue distribution of paclitaxel in CD2F1 mice. Cancer Chemother Pharmacol. 1994;34(6):465–71.

38. Seo C, Hwang YH, Kim Y, Joo BS, Yee ST, Kim CM, et al. Metabolomic study of aging in mouse plasma by gas chromatography-mass spectrometry. J Chromatogr B Anal Technol Biomed Life Sci [Internet]. 2016;1025:1–6. Available from: 10.1016/j.jchromb.2016.04.052

39. Houtkooper RH, Argmann C, Houten SM, Cant o C, Jeninga EH, Andreux eńelope A., et al. The metabolic footprint of aging in mice. Sci Rep. 2011;1:1–11.

40. Mora S, Mann G, Adegoke OAJ. Sex differences in cachexia and branched-chain amino acid metabolism following chemotherapy in mice. Physiol Rep. 2024;12(8):1–14.

41. Fujiwara T, Hattori A, Ito T, Funatsu T, Tsunoda M. Analysis of intracellular α-keto acids by HPLC with fluorescence detection. Anal Methods. 2020;12(20):2555–9.

42. Goto M, Miyahara I, Hirotsu K, Conway M, Yennawar N, Islam MM, et al. Structural determinants for branched-chain aminotransferase isozyme-specific inhibition by the anticonvulsant drug gabapentin. J Biol Chem. 2005;280(44):37246–56.

43. Wang TJ, Larson MG, Vasan RS, Cheng S, Rhee EP, McCabe E, et al. Metabolite profiles and the risk of developing diabetes. Nat Med. 2011;17(4):448–53.

44. Wurtz P, Soininen P, Kangas AJ, Rönnemaa T, Lehtimäki T, Kähönen M, et al. Branched-chain and aromatic amino acidsare predictors of insulinresistance in young adults. Diabetes Care. 2013;36(3):648–55.

45. Suryawan A, Hawes JW, Harris RA, Shimomura Y, Jenkins AE, Hutson SM. A molecular model of human branched-chain amino acid metabolism. Am J Clin Nutr. 1998;68(1):72–81.

46. Chistiakov DA, Sobenin IA, Revin V V., Orekhov AN, Bobryshev Y V. Mitochondrial aging and age-related dysfunction of mitochondria. Biomed Res Int. 2014;2014:1–7.

47. Tsuji S, Koyama S, Taniguchi R, Fujiwara T, Fujiwara H, Sato Y. Association of Serum Amino Acid Concentration With Loss of Skeletal Muscle Mass After 1 Year in Cardiac Rehabilitation Center Patients. Circ Reports. 2019;1(10):456–61.

48. Herman MA, She P, Peroni OD, Lynch CJ, Kahn BB. Adipose tissue Branched Chain Amino Acid (BCAA) metabolism modulates circulating BCAA levels. J Biol Chem. 2010;285(15):11348–56.

49. Karwi QG, Lopaschuk GD. Branched-Chain Amino Acid Metabolism in the Failing Heart. Cardiovasc Drugs Ther. 2023;37(2):413–20.

50. Uddin GM, Zhang L, Shah S, Fukushima A, Wagg CS, Gopal K, et al. Impaired Branched Chain Amino Acid Oxidation Contributes to Cardiac Insulin Resistance in Heart Failure. Cardiovasc Diabetol [Internet]. 2019;1–12. Available from: 10.1186/s12933-019-0892-3

51. Fillmore N, Wagg CS, Zhang L, Fukushima A, Lopaschuk GD. Cardiac branched-chain amino acid oxidation is reduced during insulin resistance in the heart. Am J Physiol - Endocrinol Metab. 2018;315(5):E1046–52.

52. Mullard A. Anti-ageing pipeline starts to mature. Nat Rev Drug Discov. 2018;17:609–12.

53. Hall SS. A trial for the ages. Science (80- ). 2015;349(6254):1274–8.

54. Mallappallil M, Friedman EA, Delano BG, Mcfarlane SI, Salifu MO. Chronic kidney disease in the elderly: Evaluation and management. Clin Pract. 2014;11(5):525–35.

55. Kim IH, Kisseleva T, Brenner DA. Aging and liver disease. Curr Opin Gastroenterol. 2015;31(3):184–91.

56. Vanweert F, Schrauwen P, Phielix E. Role of branched-chain amino acid metabolism in the pathogenesis of obesity and type 2 diabetes-related metabolic disturbances BCAA metabolism in type 2 diabetes. Nutr Diabetes. 2022;12(1):35.

57. Chaleckis R, Murakami I, Takada J, Kondoh H, Yanagida M. Individual variability in human blood Metabolites identifies age-related differences. Proc Natl Acad Sci U S A. 2016;113(16):4252–9.

58. Kouchiwa T, Wada K, Uchiyama M, Kasezawa N, Niisato M, Murakami H, et al. Age-related changes in serum amino acids concentrations in healthy individuals. Clin Chem Lab Med. 2012;50(5):861–70.

59. Pitkänen HT, Oja SS, Kemppainen K, Seppä JM, Mero AA. Serum amino acid concentrations in aging men and women. Amino Acids. 2003;24(4):413–21.

60. Wysokiński A, Sobów T, Kłoszewska I, Kostka T. Mechanisms of the anorexia of aging—a review. Age (Omaha). 2015;37(4).

61. Le Couteur DG, Ribeiro R, Senior A, Hsu B, Hirani V, Blyth FM, et al. Branched chain amino acids, cardiometabolic risk factors and outcomes in older men: The concord health and ageing in men project. Journals Gerontol - Ser A Biol Sci Med Sci. 2020;75(10):1805–10.

62. Imai D, Nakanishi N, Shinagawa N, Yamamoto S, Ichikawa T, Sumi M, et al. Association of Elevated Serum Branched-chain Amino Acid Levels With Longitudinal Skeletal Muscle Loss. J Endocr Soc [Internet]. 2024;8(2):1–9. Available from: 10.1210/jendso/bvad178

63. Holeček M. Branched-chain amino acids in health and disease: Metabolism, alterations in blood plasma, and as supplements. Nutrition and Metabolism. 2018.

64. Cuthbertson D, Smith K, Babraj J, Leese G, Waddell T, Atherton P, et al. Anabolic signaling deficits underlie amino acid resistance of wasting, aging muscle. FASEB J. 2005;19(3):1–20.

65. Volpi E, Sheffield-Moore M, Rasmussen BB, Wolfe RR. Basal muscle amino acid kinetics and protein synthesis in healthy young and older men. JAMA. 2001;286(10):1206–12.

66. Volpi E, Mittendorfer B, Wolf SE, Wolfe RR. Oral amino acids stimulate muscle protein anabolism in the elderly despite higher first-pass splanchnic extraction. Am J Physiol - Endocrinol Metab. 1999;277(3 40-3):513–20.

67. Balagopal P, Rooyackers OE, Adey DB, Ades PA, Nair KS. Effects of aging on in vivo synthesis of skeletal muscle myosin heavy-chain and sarcoplasmic protein in humans. Am J Physiol - Endocrinol Metab. 1997;273(4):E790–800.

68. Welle S, Thornton C, Jozefowicz R, Statt M. Myofibrillar protein synthesis in young and old men. Am J Physiol - Endocrinol Metab. 1993;264:E693–8.

69. Raue U, Slivka D, Jemiolo B, Hollon C, Trappe S. Proteolytic gene expression differs at rest and after resistance exercise between young and old women. Journals Gerontol - Ser A Biol Sci Med Sci. 2007;62(12):1407–12.

70. Edström E, Altun M, Hägglund M, Ulfhake B. Atrogin-1/MAFbx and MuRF1 are downregulated in aging-related loss of skeletal muscle. Journals Gerontol - Ser A Biol Sci Med Sci. 2006;61(7):663–74.

71. Welle S, Brooks AI, Delehanty JM, Needler N, Thornton CA. Gene expression profile of aging in human muscle. Physiol Genomics. 2003;14:149–59.

72. Ferrucci L, Fabbri E. Inflammageing: chronic inflammation in ageing, cardiovascular disease, and frailty. Physiol Behav. 2018;15(9):505–22.

73. Petry ÉR, Dresch D de F, Carvalho C, Medeiros PC, Rosa TG, de Oliveira CM, et al. Oral glutamine supplementation attenuates inflammation and oxidative stress-mediated skeletal muscle protein content degradation in immobilized rats: Role of 70 kDa heat shock protein. Free Radic Biol Med [Internet]. 2019;145(September):87–102. Available from: 10.1016/j.freeradbiomed.2019.08.033

74. Rogeri PS, Gasparini SO, Martins GL, Costa LKF, Araujo CC, Lugaresi R, et al. Crosstalk Between Skeletal Muscle and Immune System: Which Roles Do IL-6 and Glutamine Play? Front Physiol. 2020;11(October):1–11.

75. Kauppila TES, Kauppila JHK, Larsson NG. Mammalian Mitochondria and Aging: An Update. Cell Metab. 2017;25(1):57–71.

76. Sun N, Youle RJ, Fin T. The Mitochondrial Basis of Aging. Mol Cell. 2016;61(5):654– 666.

77. Spendiff S, Vuda M, Gouspillou G, Aare S, Perez A, Morais JA, et al. Denervation drives mitochondrial dysfunction in skeletal muscle of octogenarians. J Physiol. 2016;594(24):7361–79.

78. Picard M, Ritchie D, Thomas MM, Wright KJ, Hepple RT. Alterations in intrinsic mitochondrial function with aging are fiber type-specific and do not explain differential atrophy between muscles. Aging Cell. 2011;10(6):1047–55.

79. Mathieu-Costello O, Ju Y, Trejo-Morales M, Cui L. Greater capillary-fiber interface per fiber mitochondrial volume in skeletal muscles of old rats. J Appl Physiol. 2005;99(1):281–9.

80. Shen QM, Wang J, Li ZY, Tuo JY, Tan YT, Li HL, et al. Sex-Specific Correlation Analysis of Branched-Chain Amino Acids in Dietary Intakes and Plasma among Chinese Adults. J Nutr [Internet]. 2023;153(9):2709–16. Available from: 10.1016/j.tjnut.2023.07.011

81. Hernandez N, Lokhnygina Y, Ramaker ME, Ilkayeva O, Muehlbauer MJ, Crawford ML, et al. Sex Differences in Branched-chain Amino Acid and Tryptophan Metabolism and Pathogenesis of Youth-onset Type 2 Diabetes. J Clin Endocrinol Metab. 2024;109(4):e1345–58.

82. Henderson GC, Dhatariya K, Ford GC, Klaus KA, Basu R, Rizza RA, et al. Higher muscle protein synthesis in women than men across the lifespan, and failure of androgen administration to amend age related decrements. FASEB J. 2009;23(2):631–41.

83. Smith GI, Atherton P, Villareal DT, Frimel TN, Rankin D, Rennie MJ, et al. Differences in muscle protein synthesis and anabolic signaling in the postabsorptive state and in response to food in 65-80 year old men and women. PLoS One. 2008;3(3).

84. Smith GI, Atherton P, Reeds DN, Mohammed BS, Jaffery H, Rankin D, et al. No major sex differences in muscle protein synthesis rates in the postabsorptive state and during hyperinsulinemia-hyperaminoacidemia in middle-aged adults. J Appl Physiol. 2009;107(4):1308–1315.

85. Cowie CC, Rust KF, Ford ES, Eberhardt MS, Byrd-Holt DD, Li C, et al. Full accounting of diabetes and pre-diabetes in the U.S. population in 1988-1994 and 2005-2006. Diabetes Care. 2009;32(2):287–94.

86. Short KR, Vittone JL, Bigelow ML, Proctor DN, Rizza RA, Coenen-Schimke JM, et al. Impact of aerobic exercise training on age-related changes in insulin sensitivity and muscle oxidative capacity. Diabetes. 2003;52(8):1888–96.

87. Lefort N, Glancy B, Bowen B, Willis WT, Bailowitz Z, De Filippis EA, et al. Increased reactive oxygen species production and lower abundance of complex I subunits and carnitine palmitoyltransferase 1B protein despite normal mitochondrial respiration in insulin-resistant human skeletal muscle. Diabetes. 2010;59:2444–52.

88. Shin AC, Fasshauer M, Zielinski E, Lindtner C, Scherer T, O’Hare J, et al. Brain insulin regulates systemic branch-chain amino acid levels. Diabetes. 2012;20:898–909.

89. Flurkey K, Currer JM, Harrison DE. Mouse Models in Aging Research. In: The Mouse in Biomedical Research. 2007. p. 637–672.

90. Volpi E, Nazemi R, Fujita S. Muscle tissue changes with aging. Curr Opin Clin Nutr Metab Care. 2004;7(4):405–10.

91. Sonjak V, Jacob KJ, Spendiff S, Vuda M, Perez A, Miguez K, et al. Reduced Mitochondrial Content, Elevated Reactive Oxygen Species, and Modulation by Denervation in Skeletal Muscle of Prefrail or Frail Elderly Women. Journals Gerontol - Ser A Biol Sci Med Sci. 2019;74(12):1887–95.

