## Supplemental Fig 1-3 for "Effect of Age and Sex on BCAA Metabolism in Mice"

S1 Fig

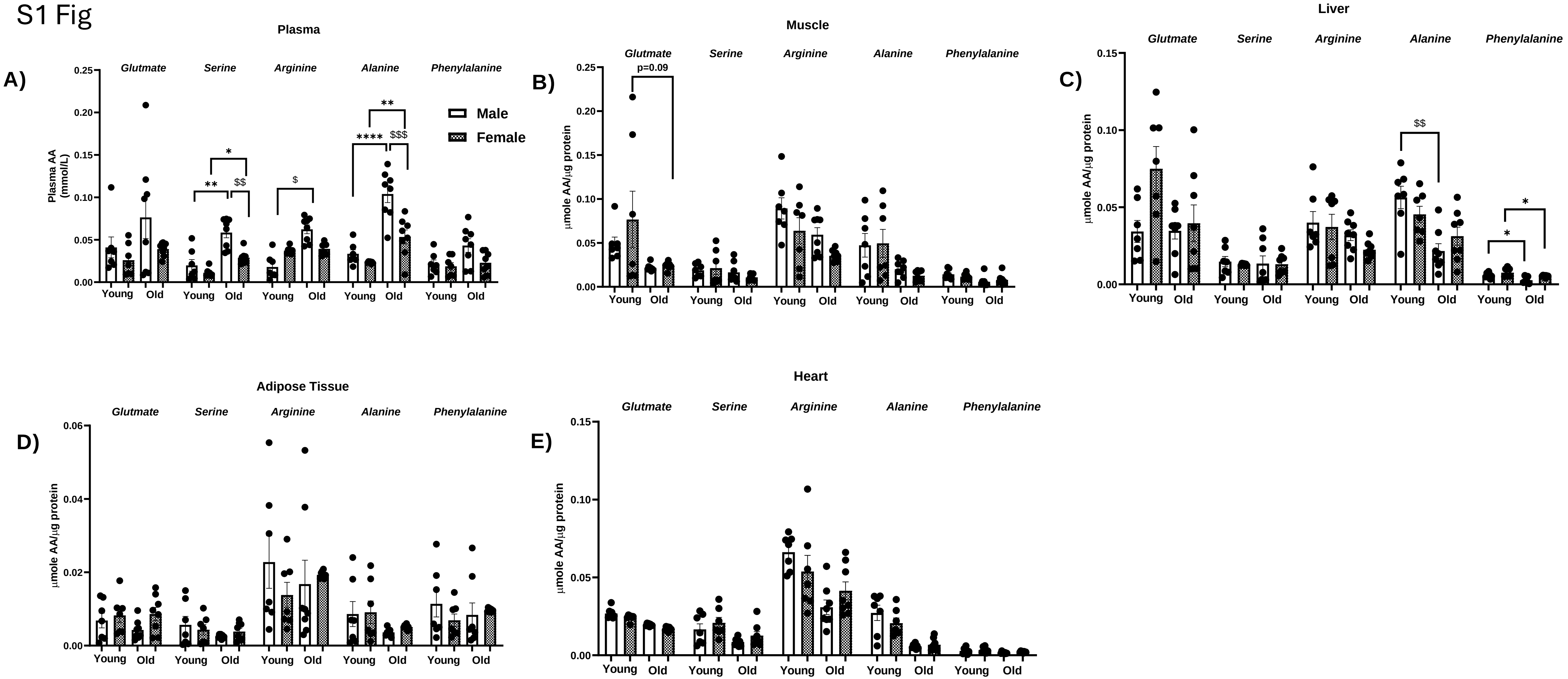
Plasma
A)
Plasma AA (mmol/L)
Glutmate
Serine
Arginine
Alanine
Phenylalanine
Male
Female
Young
Old
Muscle
B)
μmole AA/μg protein
p=0.09
Liver
C)
Adipose Tissue
D)
Heart
E)

S2 Fig

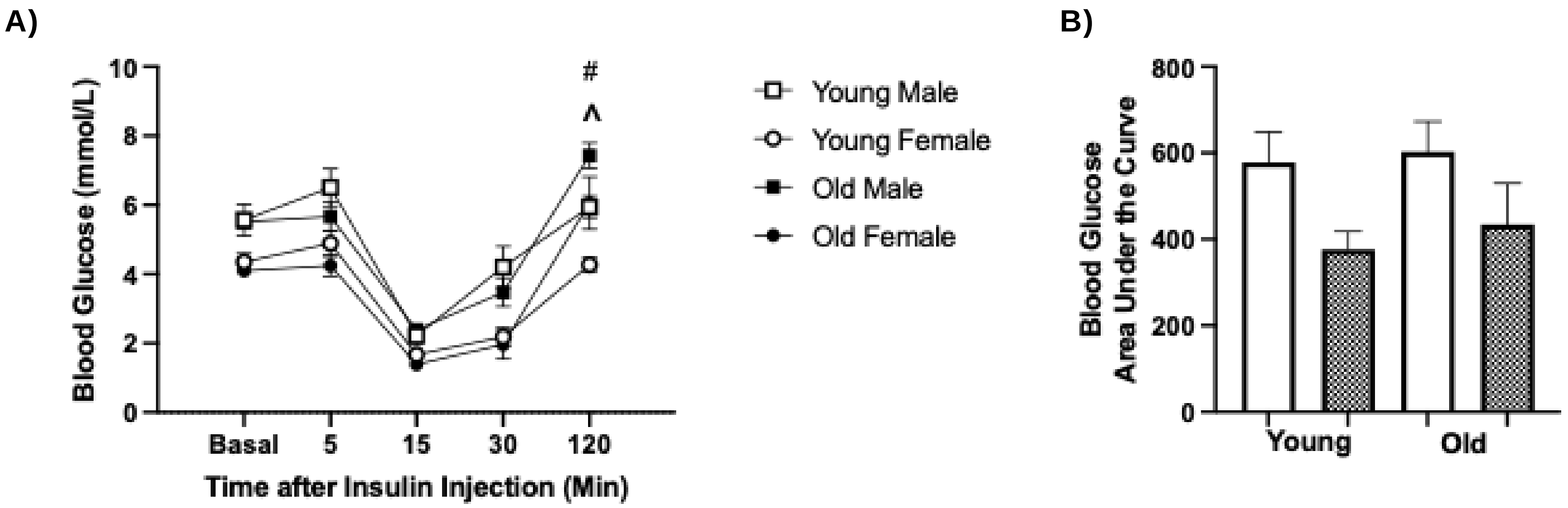
A)
Blood Glucose (mmol/L)
10
8
6
4
2
0
#
^
Basal
5
15
30
120
Time after Insulin Injection (Min)
Young Male
Young Female
Old Male
Old Female
B)
Blood Glucose
Area Under the Curve
800
600
400
200
0
Young
Old

S3 Fig

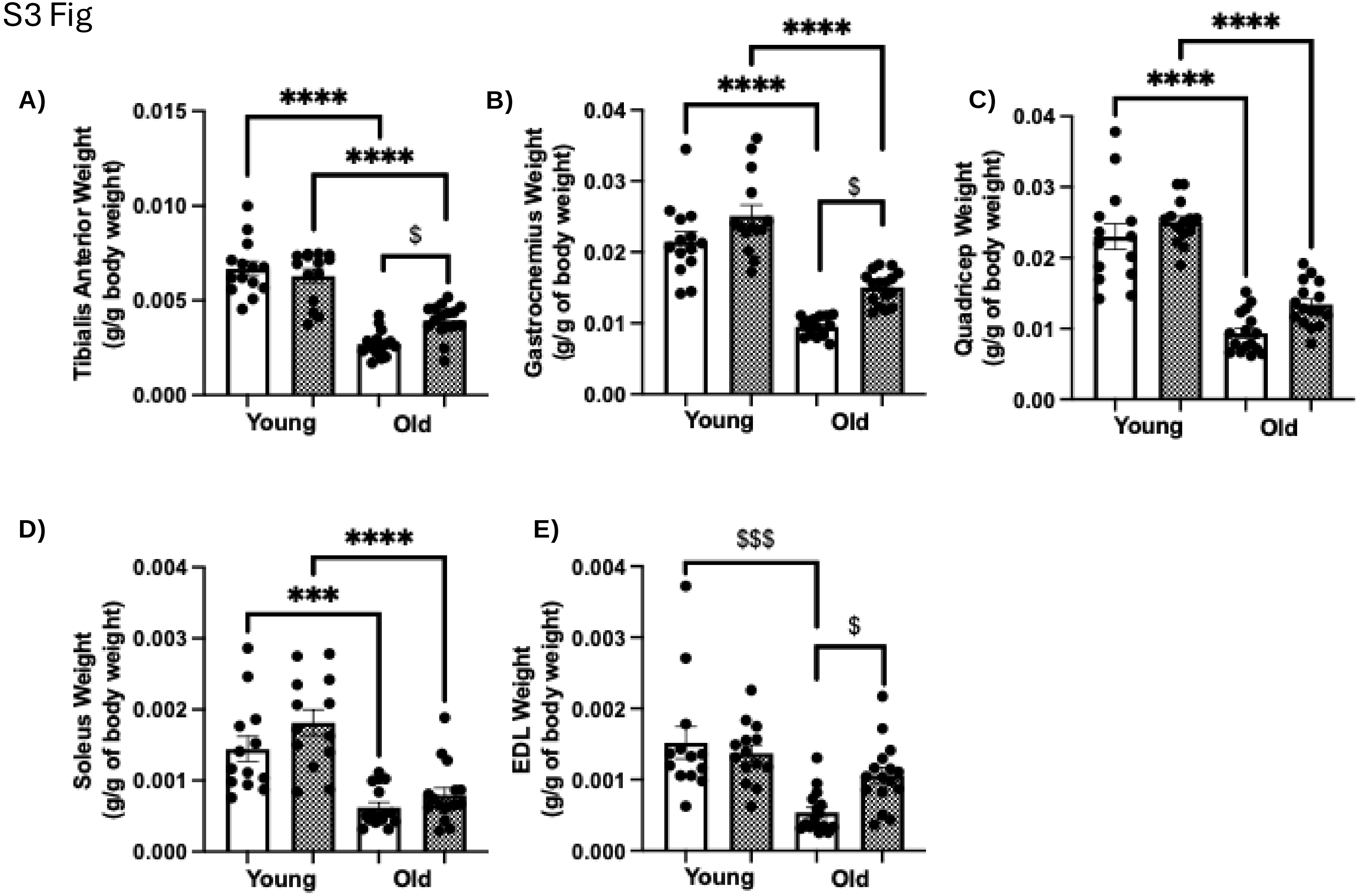
A)
Tibialis Anterior Weight
(g/g body weight)
0.015
0.010
0.005
0.000
****
****
$
Young
Old
B)
Gastrocnemius Weight
(g/g of body weight)
0.04
0.03
0.02
0.01
0.00
****
****
$
Young
Old
C)
Quadricep Weight
(g/g of body weight)
0.04
0.03
0.02
0.01
0.00
****
****
Young
Old
D)
Soleus Weight
(g/g of body weight)
0.004
0.003
0.002
0.001
0.000
****
***
Young
Old
E)
EDL Weight
(g/g of body weight)
0.004
0.003
0.002
0.001
0.000
$$$
$
Young
Old

S4 Fig

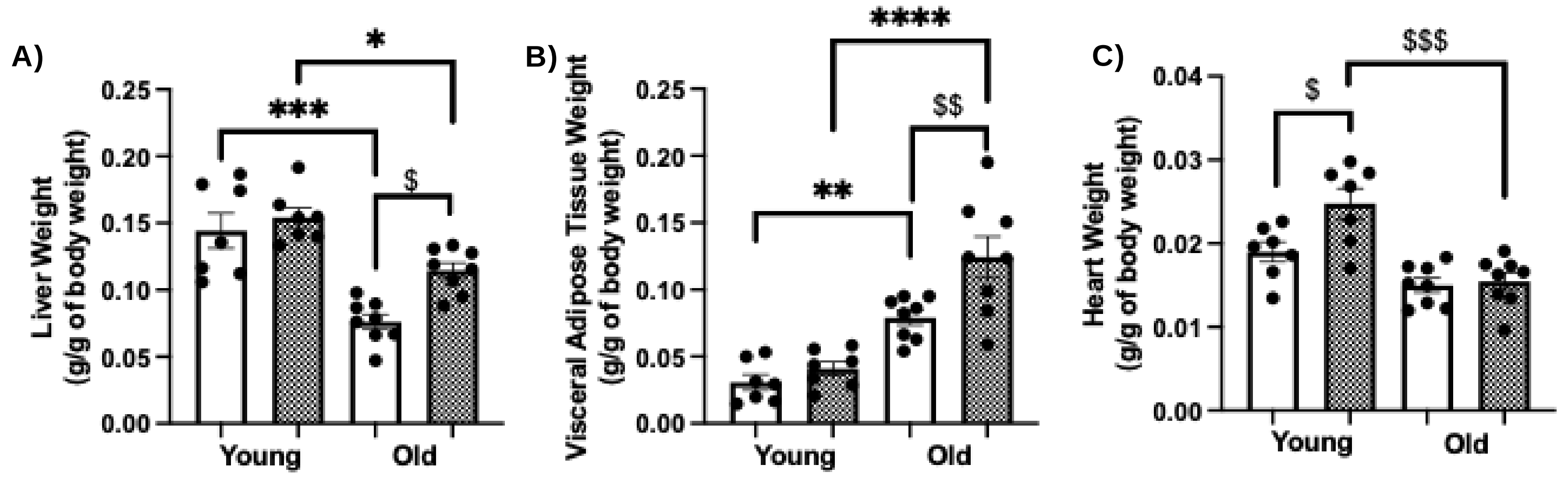
A)
Liver Weight
(g/g of body weight)
0.25
0.20
0.15
0.10
0.05
0.00
*
***
$
Young
Old
B)
Visceral Adipose Tissue Weight
(g/g of body weight)
****
$$
**
Young
Old
C)
Heart Weight
(g/g of body weight)
0.04
0.03
0.02
0.01
0.00
$
$$$
Young
Old
